# Mitragynine reduces symptoms of alcohol use disorder (and tinnitus) in a self-medication of rodents

**DOI:** 10.64898/2026.11.19.767795

**Authors:** Liubov S. Kalinichenko, Konstantin Tziridis, Nassri Abdoush, Marie-Lou Boyer, Laura Emrich, Anna Fejtova, Natasha Foerst, Renato Frischknecht, Anna Glotz, Fabian Graßl, Bernadette Hoermann, Gian-Marco Kersten, Melissa Kienberger, Salvador Kirmsse, Lisa Perschler, Lydia Scheidig, Martin Strotmann, Lissy Windhösel, Kasia Radwanska, Markus R. Heinrich, Johannes Kornhuber, Darshan Singh, Nelson Chear Jeng Yeou, Zurina Hassan, Holger Schulze, Christian P. Müller

## Abstract

Alcohol use disorder (AUD) is a major psychiatric disease. Available pharmacotherapies show rather modest compliance and efficacy. Tinnitus is a highly aversive hearing problem, that may arise from the same unsolved problems as AUD. Based on a patient report of Kratom, a Southeast Asian plant preparation, efficacy in the self-medication of AUD and tinnitus, we tested the main active Kratom alkaloid, mitragynine, for its therapeutic action in AUD mouse models covering various symptoms of the disease. We found that experimenter administered mitragynine (i.p.) inhibits well-established ongoing alcohol consumption in a social setting when alcohol is permanently available. Mitragynine also inhibits alcohol consumption in a non-social, isolated setting. While mitragynine did not work for prophylaxis during abstinence, it markedly attenuated the relapse to binge drinking with alcohol availability restricted to a limited time. Furthermore, mitragynine reduced alcohol seeking behaviour during its establishment as well as during retrieval from memory. It was also effective to attenuate its re-establishment after extinction. We found significant antidepressive effects of mitragynine, during alcohol withdrawal. To address the compliance problem for pharmacotherapies, we established a test that can monitor this and found that mice after a short training, learned to orally self-administer mitragynine hydrochloride leading to a strong reduction in voluntary alcohol drinking. In parallel, mitragynine reduced tinnitus after a noise trauma in a gerbil model. Together, these findings suggest that mitragynine may be used as a pharmacotherapy for AUD and tinnitus with an expected high compliance rate.

## Introduction

Alcohol use disorder (AUD) is a major psychiatric disorder resulting from loss of control over alcohol consumption (Carvalho et al., 2019). Alcohol consumption is a nearly worldwide phenomenon, as shown by the majority of adult people in many cultures around the world (Wadley, 2016). Thereby, alcohol is not only consumed for hedonic reasons, but often instrumentalized to self-manage various tasks and problems. Among them are mood disorders, such as anxiety and depression, and related physical symptoms (Müller et al., 2001, 2023). However, recent studies suggest that, for overall health, there is no safe level or pattern of alcohol consumption in humans (Rehm et al., 2017; GBD, 2020; Shield et al., 2020). AUD is a highly prevalent disorder with over 600 million people currently suffering from AUD or alcohol dependence (WHO, 2024).

There is a strong need for better therapies for AUD and related symptoms. While there are prescribable pharmacotherapies available (Fairbanks et al., 2020; Dionisi et al., 2024), they are often not used by patients due to low efficacy and insufficient compliance (McPheeters et al., 2023; Kotake et al., 2024; Wellensiek et al., 2024). As such, not only efficacy, but also compliant use of a new candidate drug needs to be assured in the search of a new pharmacotherapeutic compound as a criterion in drug development.

While there are a plethora of novel compounds generated and tested to reduce AUD symptoms based on a neuropharmacological treatment rationale (Walker and Lawrence, 2018; Koob, 2024), other ways of searching may comprise those substances that patients have identified for their own self-management of AUD on a trial-and-error basis with already available compounds. We recently described a patient who reported a long-term successful self-medication of depressive symptoms with the plant preparation of Kratom (Müller et al., 2021), a Southeast Asian medical plant (Hassan et al., 2013). In the course of this self-medication, he also admitted to having completely abstained from alcohol, for which he had a verified AUD diagnosis, and his tinnitus also came under control (Müller et al., 2021). Kratom is in many countries a freely available drug used as a food supplement with a long history of medical use for several symptoms in Southeast Asia (Hassan et al., 2013; Hartley et al., 2022). Its main psychoactive alkaloid is the highly abundant mitragynine (Flores-Bocanegra et al., 2020; McCurdy et al., 2024).

In rodents and humans, tinnitus is a symptom of hearing loss, independent whether it is clinically relevant or “hidden” (Schaette & McAlpine, 2011; Tziridis et al, 2021 ; Mazurek et al., 2022). It affects between 12 and 30% of human adults and constitutes a substantial clinical and socioeconomic burden (McCormack et al., 2016; Tziridis et al., 2022). Although often transient, a significant subset of patients develops chronic and distressing tinnitus accompanied by sleep disruption, cognitive impairment, and affective comorbidities, including anxiety and depression (Langguth et al., 2013). For these patients, the standard therapy is usually counselling and behavioural therapy (Mazurek et al., 2022), as many patients show low compliance in using hearing aids to treat any hearing loss. Currently, there are no effective pharmacotherapies for tinnitus available.

We have taken this report as a reason to investigate whether the main psychoactive compound of Kratom, mitragynine, may indeed inhibit AUD symptoms at a large scale and, in a parallel approach, also reduce tinnitus in preclinical approaches. As compliance is a major problem for new AUD therapies, we also tested for the first time whether this compound is voluntarily consumed as a self-medication that, in parallel, reduces alcohol consumption.

## Results

### Mitragynine inhibits social alcohol drinking

A common scenario of alcohol consumption is social drinking, when individuals are in a social environment and alcohol is only available for some time of the day (MacKillop et al., 2022). We tested the effects of mitragynine on social drinking in male mice after a stable 20 vol. % alcohol consumption was gradually established and maintained for four weeks with intermittent access. Mitragynine treatment (5 mg/kg/day, i.p.) significantly reduced established alcohol drinking in these mice during the three treatment days, but not thereafter (Fig. 1A). Mitragynine had no effect on water consumption during the whole time of measurement (Fig. 1B).

**Figure 1.**
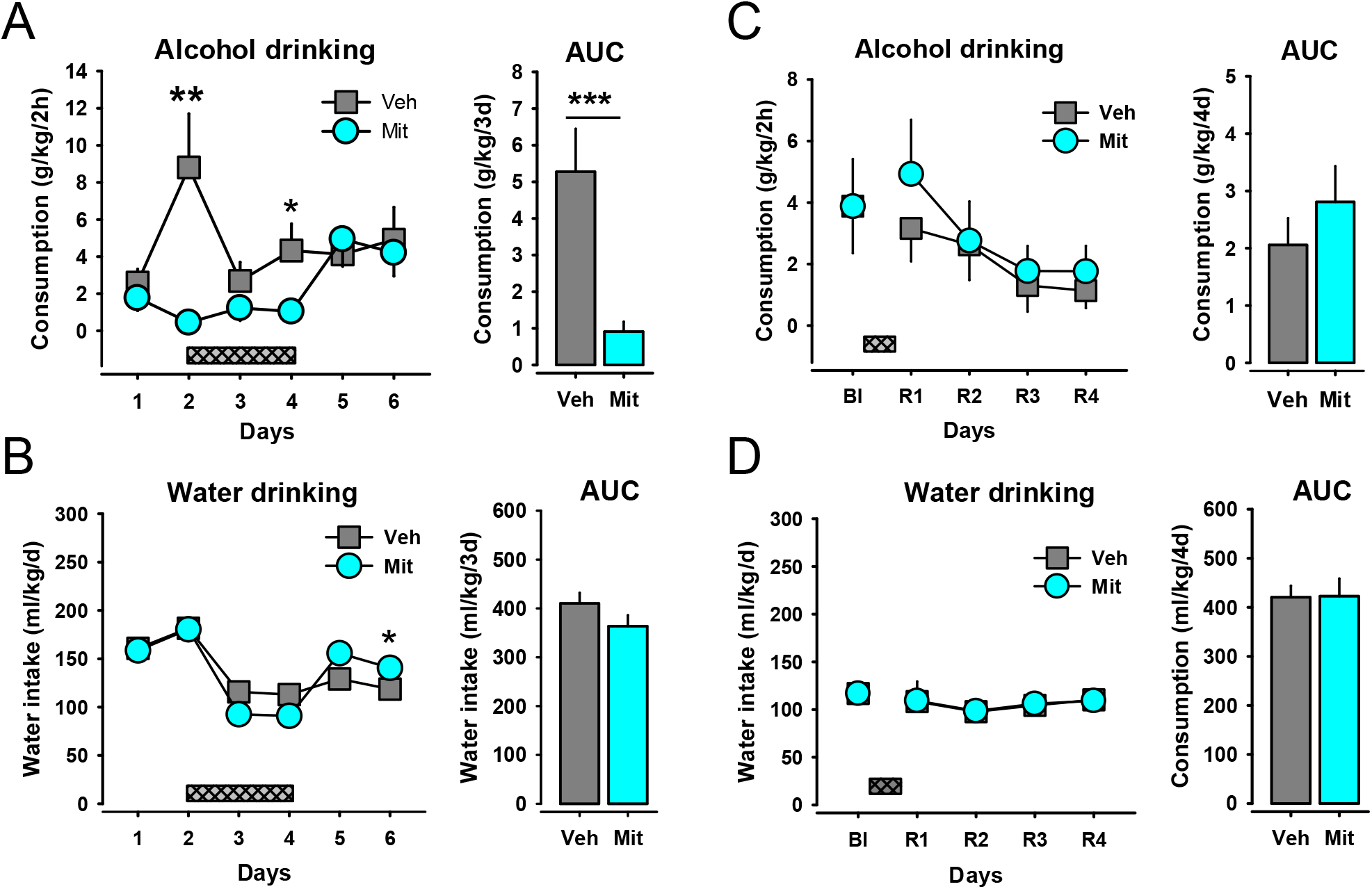
Mitragynine inhibits well-established alcohol drinking in a social setting. **A** Alcohol consumption in an IntelliCage of mice during mitragynine treatment on three days (5 mg/kg, i.p., grey bar). A two-way ANOVA suggested an effect of treatment (F_1,26_=4.106, p=0.05), time (F_5,130_=2.319, p=0.0469), and interaction (F_5,130_=3.742, p=0.003). Area under the curve (AUC) analyses of average drinking only for the treatment days confirmed a significant effect of mitragynine (t(41)=-3.567, p<0.001). **B** Effect of mitragynine on water consumption. A two-way ANOVA suggested no effect of treatment (F_1,26_<0.001, p=0.998), but an effect of time (F_5,130_=28.024, p<0.001), and interaction (F_5,130_=3.204, p=0.009). AUC analyses of average water drinking only for the treatment days confirmed a non-significant trend effect of mitragynine (t(41)=-2.0141, p=0.051). **C.** Mitragynine prophylaxis (grey bar) had no effect on alcohol drinking (treatment: F_1,26_=0.180, p=0.674), time: F_4,104_=5.786, p<0.001, interaction: F_4,104_=0.4629, p=0.763; AUC: t(55)=-1.258, p=0.214), or **D.** water consumption during an alcohol deprivation effect (ADE; treatment: F_1,26_=0.002, p=0.963, time: F_4,104_=1.994, p=0.101, interaction: F_5,104_=0.011, p=0.999; AUC: t(55)=-0.079, p=0.937), when given on 3 days during withdrawal immediately before reinstatement of drinking. Data are expressed as means ± s.e.m.; *p<0.05; **p<0.01; ***p<0.001 vs. vehicle).

In a second test, we probed whether mitragynine would work as a prophylaxis during periods of abstinence that precede a relapse. Therefore, we tested the same animals again after they underwent a three-week withdrawal period. Mitragynine was administered three days before alcohol re-exposure and the alcohol deprivation effect (ADE), a short overshoot of consumption after abstinence (Spanagel and Hölter, 2000), was measured. We found that mitragynine did not affect the ADE expression or water consumption on that occasion (Fig. 1C-D). These data suggest that mitragynine inhibits well-established alcohol drinking in a social setting.

### Mitragynine inhibits non-social alcohol drinking, but not relapse

Another typical drinking scenario is isolated alcohol drinking with near-unlimited alcohol availability. We tested the effects of mitragynine on isolated alcohol drinking in male mice after a stable 16 vol.% alcohol consumption was gradually established and maintained for two weeks with unlimited access in a two-bottle-free choice paradigm. Mitragynine treatment significantly reduced established alcohol drinking in these mice during the treatment days, but not thereafter (Fig. 2A). Mitragynine had a small effect on alcohol preference vs water (Fig. 2B), but did not affect water consumption during the whole time of measurement (Fig.2C).

**Figure 2.**
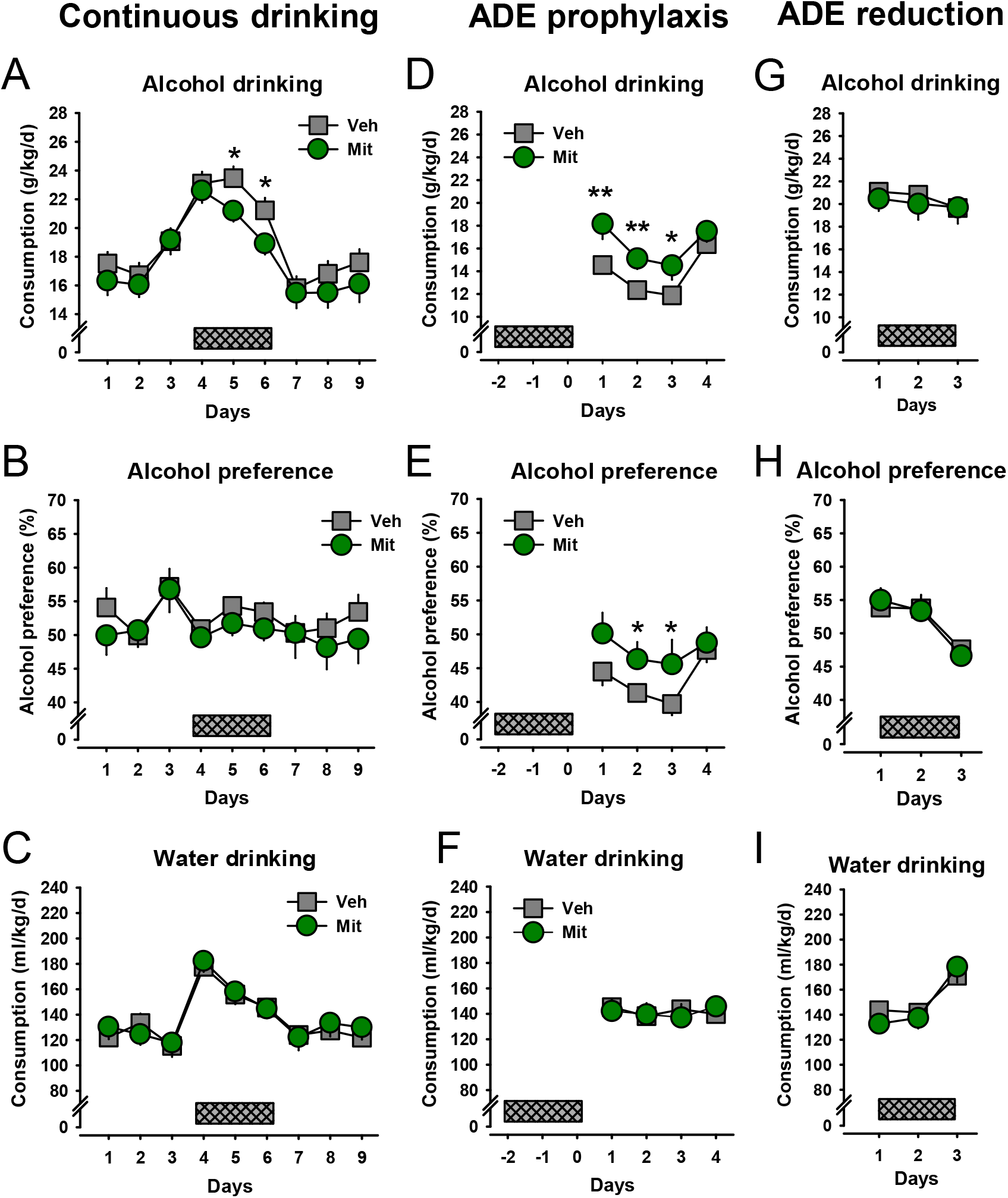
Mitragynine inhibits well-established free-access alcohol drinking in a non-social setting. **A.** Alcohol consumption in a two-bottle free-choice test of male mice during mitragynine treatment on three days (5 mg/kg, i.p., grey bar; two-way ANOVA; treatment: F_1,29_=1.193, p=0.284, time: F_8,232_=28.036, p<0.001; interaction: F_8,232_=4.785, p=0.724). **B.** Effect of mitragynine on alcohol preference vs. water (treatment: F_1,29_=0.465, p=0.501, time: F_8,232_=2.838, p=0.005, interaction: F_8,232_=0.436, p=0.898) and **C.** on water consumption (treatment: F_1,29_=0.051, p=0.823, time: F_8,232_=22.832, p<0.001, interaction: F_8,232_=0.421, p=0.908). **D.** Mitragynine enhanced the alcohol deprivation effect (ADE) when given on 3 days before reinstatement (grey bar) of alcohol consumption (treatment: F_1,30_=6.243, p=0.018, time: F_3,90_=20.920, p<0.001, interaction: F_3,90_=1.579, p=0.200), and **E.** alcohol preference (treatment: F_1,30_=3.280, p<0.001, time: F_3,90_=5.634, p=0.001, interaction: F_3,90_=0.987, p=0.402), but not on **F.** water consumption (treatment: F_1,30_=0.004, p=0.950, time: F_3,90_=0.425, p=0.736, interaction: F_3,90_=0.623, p=0.602). **G.** No effect of mitragynine during the expression of an alcohol deprivation effect when given on 3 days during re-exposure to alcohol (grey bar) on alcohol drinking (treatment: F_1,25_=0.140, p=0.711, time: F_2,50_=1.825, p=0.172, interaction: F_2,50_=0.289, p=0.750), on **H.** alcohol preference (treatment: F_1,25_=0.001, p=0.975, time: F_2,50_=19.393, p<0.001, interaction: F_2,50_=0.328, p=0.722), and on **I.** water consumption (treatment: F_1,25_=0.118, p=0.734, time: F_2,50_=22.547, p<0.001, interaction: F_2,50_=1.128, p=0.332). Data are expressed as means ± s.e.m. (*p<0.05; **p<0.01 vs. vehicle).

In a second test, we probed whether mitragynine would work as a prophylaxis during periods of abstinence that precede a relapse. Therefore, we tested the same animals again after they underwent a three-week withdrawal period. Mitragynine was administered three days before alcohol re-exposure. We found that mitragynine actually enhanced relapse drinking ADE and alcohol preference, but did not affect water drinking in this situation (Fig. 2D-F).

In a third test, we asked whether mitragynine would reduce the ADE when given during re-lapse had started. Here we found no effect of mitragynine on either alcohol or water drinking and preference (Fig. 2G-I). These data suggest that mitragynine inhibits well-established isolated alcohol drinking, but may enhance relapse drinking when given during withdrawal as prophylaxis.

### Mitragynine reduces binge-drinking relapse

Alcohol binge drinking, the consumption of high amounts of alcohol in a short time, is a major problem in AUD (MacKillop et al., 2022). Here we tested whether mitragynine would be able to reduce this particular type of consumption under three different drinking scenarios. We investigated the effects of mitragynine on isolated drinking in male mice after a stable 20 vol.% alcohol consumption in a drinking-in-the-dark paradigm when alcohol is only at restricted times during the dark active period of the mice with unlimited access in a two-bottle free-choice assay. Mitragynine treatment (5 mg/kg, i.p.) did not change high-level alcohol drinking, alcohol preference or water drinking during an ongoing drinking episode (Fig. 3A-C).

**Figure 3.**
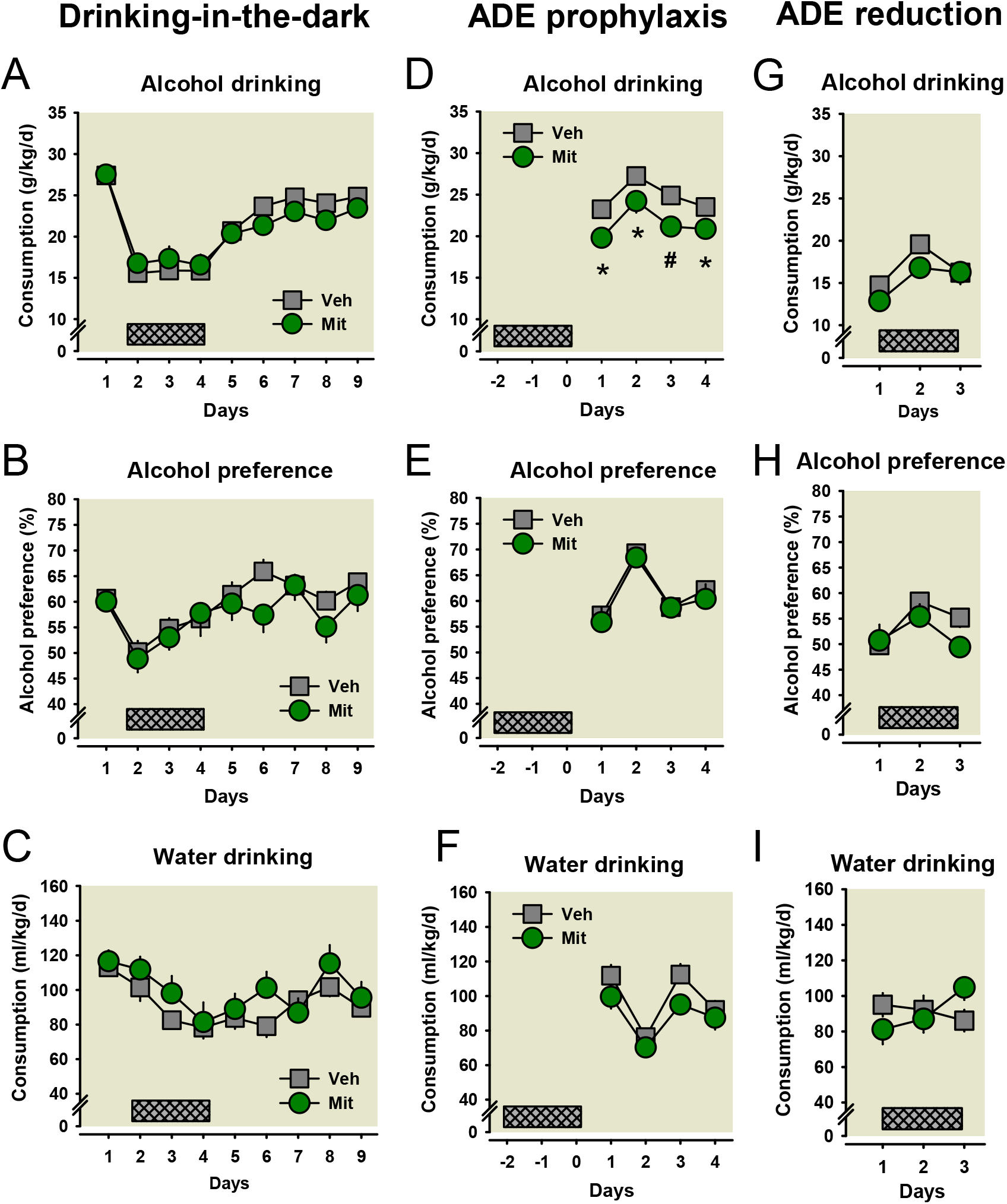
Mitragynine reduces binge-drinking relapse in a non-social setting. **A.** Alcohol consumption in a two-bottle free-choice test of mice drinking-in-the-dark during mitragynine treatment on three days (5 mg/kg, i.p.) (treatment: F_1,28_=0.823, p=0.372, time: F_8,224_=10.489, p<0.001, interaction: F_8,224_=1.840, p=0.071), **B.** alcohol preference (treatment: F_1,28_=0.383, p=0.541, time: F_8,224_=58.580, p<0.001, interaction: F_8,224_=1.096, p=0.367), and **C.** water consumption (treatment: F_1,28_=1.041, p=0.406, time: F_8,224_=7.777, p<0.001, interaction: F_8,224_=1.347, p=0.256). **D.** Mitragynine attenuated binge-like alcohol when given on 3 days before reinstatement of alcohol consumption after withdrawal (treatment: F_1,27_=11.828, p=0.002, time: F_4,108_=15.073, p<0.001, interaction: F_4,108_=0.260, p=0.903), but had no effect on **E.** alcohol preference (treatment: F_1,27_=0.390, p=0.537, time: F_4,108_=30.493, p<0.001, interaction: F_4,108_=0.291, p=0.883), and **F.** water consumption (treatment: F_1,27_=1.604, p=0.216, time: F_4,108_=20.977, p<0.001, interaction: F_4,108_=0.959, p=0.433). **G.** Mitragynine significantly reduced alcohol the relapse to binge-drinking when given during re-exposure shown as average consumption over the three treatment days (treatment: F_1,24_=5.092, p=0.033, time: F_7,168_=15.819, p<0.001, interaction: F_7,168_=1.285, p=0.261). **H.** alcohol preference (treatment: F_1,24_=0.901, p=0.352, time: F_7,168_=3.236, p=0.003, interaction: F_7,168_=1.069, p=0.386), and **I.** water consumption (treatment: F_1,24_=0.284, p=0.599, time: F_7,168_=6.861, p<0.001, interaction: F_7,168_=1.575, p=0.146). (Data are expressed as means ± s.e.m.; *p<0.05; ^#^p<0.01 vs. vehicle).

In a second test, we probed whether mitragynine would work as a prophylaxis during periods of abstinence that precede a relapse to binge-drinking. We tested the same animals again after they underwent a three-week withdrawal period. Mitragynine (5 mg/kg, i.p.) was administered three days before alcohol re-exposure. We found that mitragynine significantly reduced relapse drinking (Fig. 3D), but had no significant effect on alcohol preference or water drinking in this situation (Fig. 3E-F).

In a third test, we investigated how mitragynine (5 mg/kg, i.p.) would affect alcohol binge-drinking when administered during relapse (Fig. 3G). We found a small, but in total significant reduction of alcohol binge-drinking in this drinking scenario, but no effect on alcohol preference or water consumption (Fig. 3H-I). These data suggest that mitragynine may prophylactically reduce binge-drinking relapse, but may not affect binge-drinking while ongoing.

### Mitragynine blocks the establishment of alcohol seeking

Drug-seeking behaviour is a hallmark of AUD (MacKillop et al., 2022). It is a memory-based behaviour that often dominates behavioural repertoire in AUD (Robbins et al., 2008; Müller and Schumann, 2011). Here we tested whether mitragynine may help to avoid the establishment of drug-seeking behaviour. This was tested in the establishment of alcohol induced conditioned place preference (CPP; Huston et al., 2013). Thereby, animals were treated repeatedly with mitragynine (0, 5, 10 mg/kg; i.p.) before each CPP learning trial. We found that mitragynine dose-dependently attenuated the establishment of an alcohol CPP, as it reduced the increase of time spend in a drug-paired environment (Fig. 4A). This was caused by a reduction in time spend in this environment at each entry (Fig. 4B) and by locomotion during each entry (Fig. 4D). Conditioned hyperlocomotion that usually develops after alcohol conditioning was prevented by attenuation of the acute alcohol effects on locomotor activity in each learning trial (Fig. 4C). Altogether, these findings suggest that mitragynine can fully prevent the learning of conditioned alcohol seeking behaviour in a dose-dependent way.

**Figure 4.**
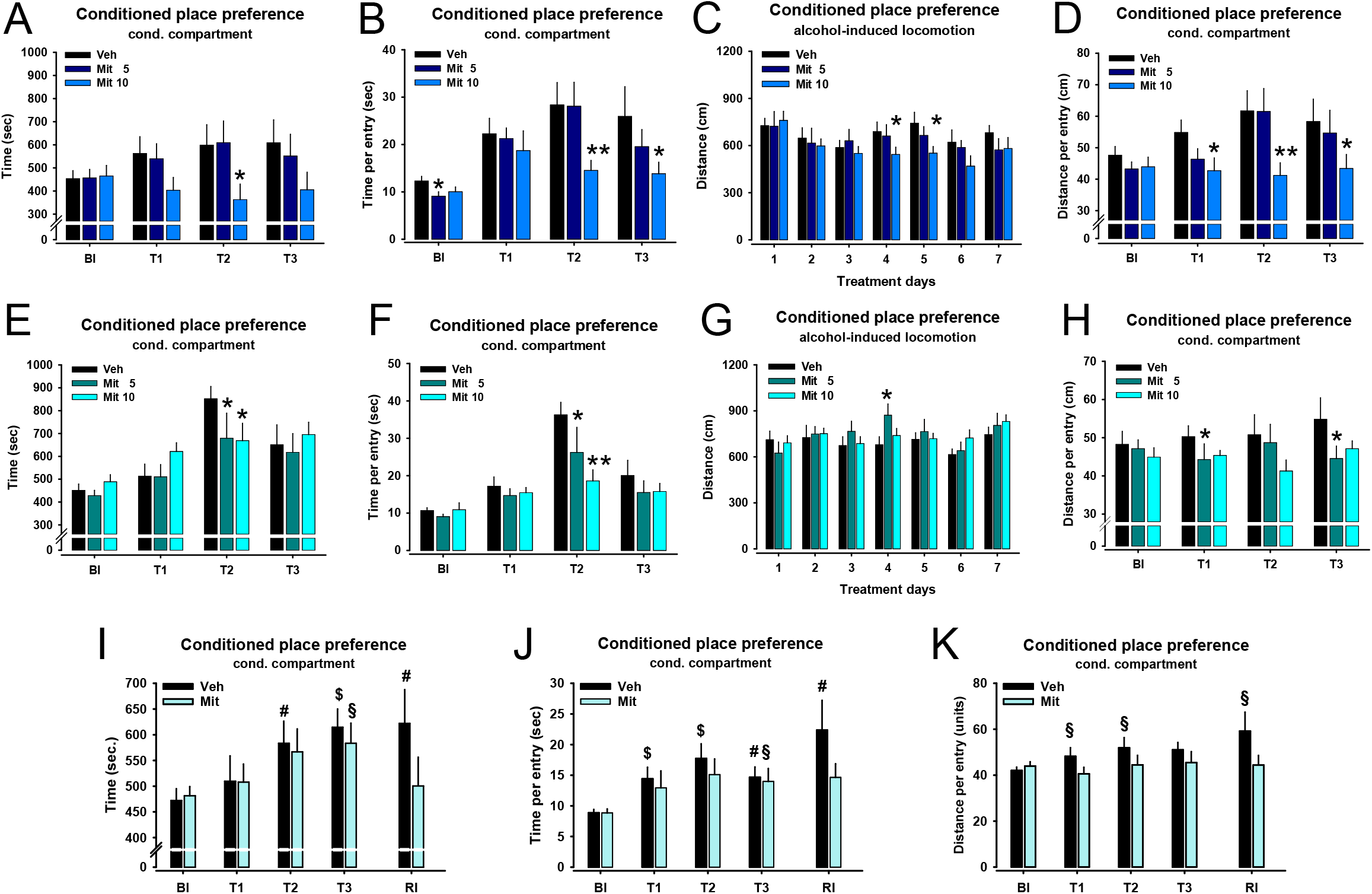
The effects of mitragynine on learning and retrieval of alcohol seeking behaviour. Effects of mitragynine (Mit, in mg/kg; i.p.) on the establishment of an alcohol-induced conditioned place preference (CPP; n=12/group). **A.** Time in the conditioning compartment (two-way ANOVA; treatment: F_2,27_=0.628, p=0.541, time: F_3,81_=3.204, p=0.028, interaction: F_6,81_=3.666, p=0.003). **B.** Time per entry in the conditioning compartment (treatment: F_2,27_=2.384, p=0.111, time: F_3,81_=11.988, p<0.001, interaction: F_3,81_=1.578, p=0.164). **C.** Distance moved during conditioning trials (treatment: F_2,28_=0.966, p=0.393, time: F_6,168_=3.489, p=0.003, interaction: F_12,168_=0.745, p=0.706), and **D.** Distance moved per entry in the conditioning compartment (treatment: F_2,27_=2.557, p=0.096, time: F_3,81_=4.518, p=0.006, interaction: F_6,81_=1.835, p=0.103). Effects of mitragynine on the retrieval of an alcohol-induced CPP (n=11-12 /group). **E.** Time in the conditioning compartment (ANOVA; treatment: F_2,27_=0.888, p=0.423, time: F_3,81_=18.500, p<0.001, interaction: F_6,81_=1.918, p=0.088). **F.** Time per entry in the conditioning compartment (treatment: F_2,27_=4.130, p=0.028, time: F_3,81_=19.673, p<0.001, interaction: F_3,81_=3.246, p=0.007), **G.** Distance moved during conditioning trials (treatment: F_2,30_=0.541, p=0.588, time: F_6,180_=2.348, p=0.033, interaction: F_12,180_=1.442, p=0.151), and **H.** Distance moved per entry in the conditioning compartment (treatment: F_2,27_=3.471, p=0.046, time: F_3,81_=0.853, p=0.469, interaction: F_6,81_=0.861, p=0.527) (Bl – baseline, T-test trial). Data are expressed as means ± s.e.m. (*p≤0.05, **p≤0.01 vs. vehicle). Effects of mitragynine on the re-instatement of an alcohol-induced CPP after extinction (n=14-15/group). **I.** Time in the conditioning compartment (treatment: F_1,19_=1.717, p=0.206, time: F_4,76_=5.298, p<0.001, interaction: F_4,76_=1.709, p=0.157). **J.** Time per entry in the conditioning compartment (treatment: F_1,19_=5.615, p=0.029, time: F_4,76_=6.626, p<0.001, interaction: F_4,76_=1.239, p=0.301), **K.** Distance moved during conditioning trials (treatment: F_1,19_=7.643, p=0.012, time: F_4,76_=2.541, p<0.001, interaction: F_4,76_=1.482, p=0.216) (Bl – baseline, T-test trial; RI – reinstatement test trial). Data are expressed as means ± s.e.m. (^§^p≤0.05, ^#^p≤0.01, ^$^p≤0.001 vs. baseline).

### Mitragynine blocks the retrieval of alcohol seeking behaviour

A key mechanism to reduce AUD symptoms in a therapy would be to reduce the retrieval of seeking behaviour and subsequent relapse (MacKillop et al., 2022). Here we tested whether mitragynine can reduce the retrieval of drug-seeking behaviour in a CPP test (König et al., 2020). Thereby, animals were treated repeatedly with mitragynine (0, 5, 10 mg/kg; i.p.) before each CPP retrieval episode. We found that mitragynine attenuated the retrieval of an alcohol CPP, but only at certain time points of the learning curve, as measured by the time spent in a drug-associated environment (Fig. 4E). This reduction in dwell time went along with a reduction in time spend in this environment at each entry (Fig. 4F) and in locomotion during each entry (Fig. 4H). These effects were evident after three learning trials, but less so after seven trials. There was no systematic effect of the treatment on alcohol-induced hyperlocomotion at each conditioning trial (Fig. 4G). Overall, these findings may suggest that mitragynine at least temporarily inhibits the retrieval of conditioned alcohol seeking behaviour.

### Mitragynine facilitates the extinction of alcohol seeking behaviour

A major goal for a pharmacotherapy of AUD would be to not only block the expression of drug-related behaviours, but ideally to extinguish them. Here we investigated whether mitragynine has the potential to facilitate the extinction of alcohol seeking behaviour in a test of alcohol CPP extinction. Male mice were trained to establish a CPP for alcohol (2 g/kg; i.p.), which was significantly expressed after three conditioning trials, and then maintained during further learning trials (Fig. 4I). Thereafter, animals received five extinction learning trials, when no alcohol was given before pairing. Then animals were dived into two groups with one receiving mitragynine and the other vehicle, preceding two re-instatement trials with alcohol (2 mg/kg; i.p.). Persistence of the CPP was tested one day later in a re-instatement test. We found that both groups established an alcohol CPP. This CPP was stable after three and seven condition trials, and persistent after re-instatement with vehicle treatment. Mitragynine (5 mg/kg; i.p.) significantly reduced the CPP in the re-instatement test (Fig. 4I). This effect was driven by a reduction in time as well as locomotion in the conditioning compartment during re-instatement test (Fig. 4J-K). Altogether, these findings may suggest that mitragynine can interfere with re-instatement of alcohol seeking behaviour after an extinction and efficiently reduce it.

### Mitragynine delays alcohol sedation effects, but extends duration

Often sought-after effects of alcohol are its relaxing and sedating action (Müller et al. 2021; MacKillop et al., 2022), which may also be a predictor for the development of AUD (Spanagel, 2009). Here we tested whether mitragynine can influence the acute sedating effects of alcohol (3.5 mg/kg; i.p.) on the one hand and their development over time of alcohol exposure on the other hand. We measured the loss of rightening reflex (LORR) in naïve male mice and found that mitragynine (0, 5, 10 mg/kg; i.p.) did not significantly affect LORR onset (Fig. 5A), but dose-dependently prolonged LORR duration (Fig. 5-B).

**Figure 5.**
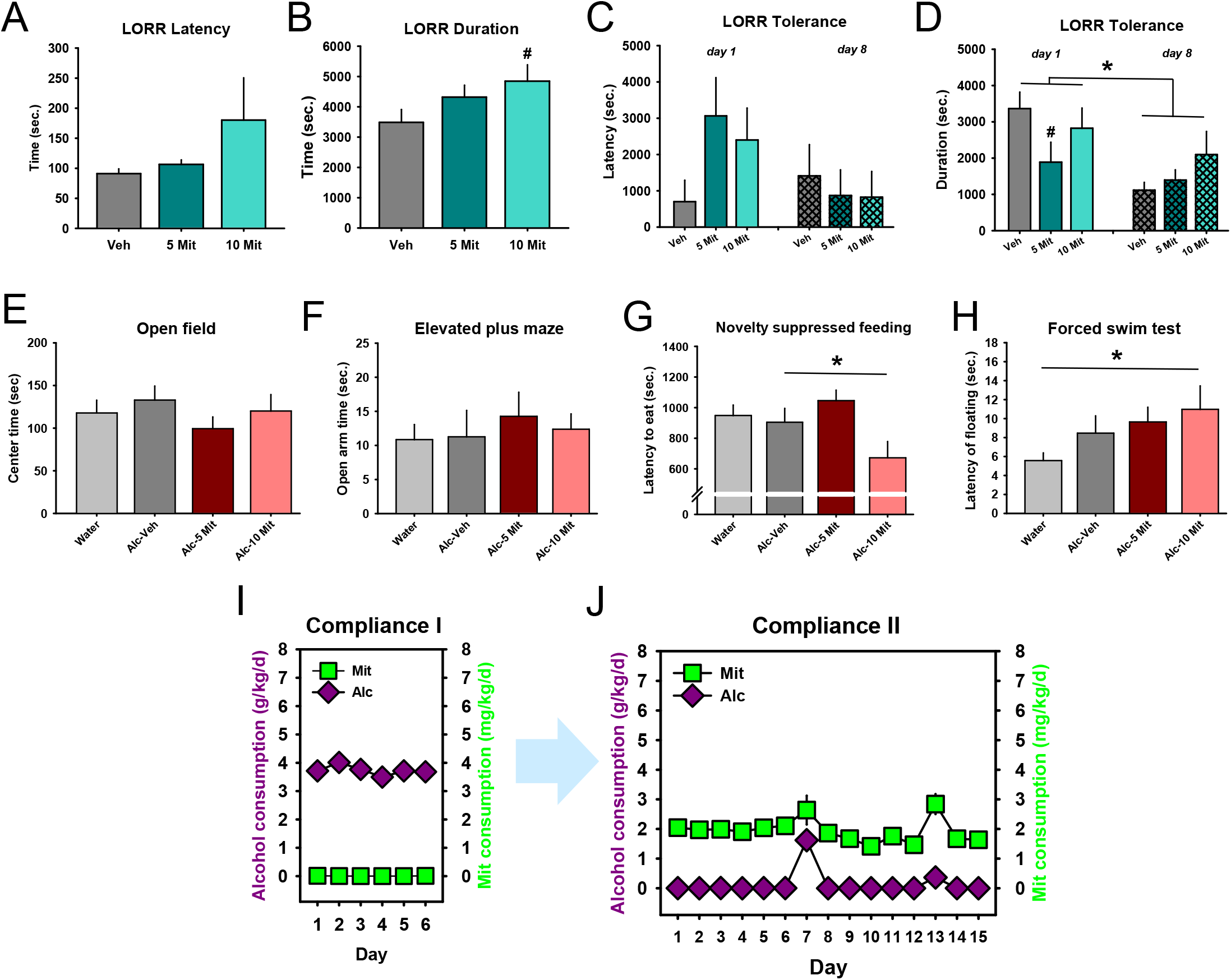
The effects of mitragynine on sedating and withdrawal effects of alcohol, and compliance in self-medication. Effects of mitragynine on the loss of rightening reflex (LORR; n=8-11/group). **A.** Latency to lose the LORR after single alcohol administration (3.5 mg/kg; i.p.) (one-way ANOVA; treatment: F_2,25_=1.723, p=0.199). **B.** Duration of the LORR (F_2,25_=2.3639, p=0.1147). **C.** Latency to lose the LORR after alcohol administration (3.5 mg/kg; i.p.) in alcohol experienced mice with tolerance development (n=10-12/group; two-way ANOVA; treatment: F_2,28_=0.6444, p=0.5326, time: F_1,28_=2.1101, p=0.1574, and interaction: F_2,28_=1.3902, p=0.2656). **D.** Duration to lose the LORR after alcohol administration in alcohol experienced mice with tolerance development (two-way ANOVA; treatment: F_2,28_=1.4364, p=0.2547, time: F_1,28_=6.5398, p=0.0163, and interaction: F_2,28_=2.2867, p=0.1203). Mitragynine effects on emotional behaviour during withdrawal from alcohol drinking. **E.** Center time in the open field test of anxiety during alcohol withdrawal vs. no alcohol drinking control (one-way ANOVA; treatment: F_2,36_=0.954, p=0.396). **F.** Open arm time in the elevated plus maze test of anxiety (F_2,37_=0.250, p=0.780). **G.** Latency to eat in the novelty suppressed feeding test of depression (F_1,40_=4.905, p=0.013). **H.** Latency of floating behaviour in the forced swim test of depression (F_1,36_=0.497, p=0.613). Data are expressed as means ± s.e.m. (*p<0.05). Alcohol drinking in mice in the social environment of the IntelliCage (n=12/cage). **I.** After four weeks of 16 vol.% alcohol consumption, access to mitragynine (Mit; p.o.; FR1) was given for 6 days. **J.** Alcohol and mitragynine p.o. self-administration (both: FR1) after one week of forced mitragynine consumption during which only mitragynine solution was available. Data are expressed as means ± s.e.m.

In other animals, that had experienced alcohol for several times (7 daily injections, 3.5 mg/kg; i.p.), LORR showed tolerance development, indicated by a shortened time of duration. Pretreatment with mitragynine (0, 5, 10 mg/kg; i.p.) did no longer prolong LORR duration (Fig. 5 C-D). These data may suggest that mitragynine may enhance one of the desired alcohol effects. This effect may, however, disappear with tolerance development after repeated alcohol exposure.

### Antidepressant mitragynine effects during alcohol withdrawal

A strong driving force for the establishment and maintenance of an AUD is the aversive effects of alcohol in withdrawal. Alcohol consumption and acute alcohol effects are often followed by a long time period of depressive mood and anhedonia, which can be immediately attenuated by new alcohol intake (Koob and Volkow, 2016). Attenuating the severity of withdrawal effects may, thus, reduce the chance of resuming consumption during withdrawal. Here we tested how mitragynine influences the emotional state during alcohol withdrawal. Naïve male mice were trained to drink alcohol (10 vol.%) in a DID paradigm for six weeks (Pang et al., 2013). Then a two-week withdrawal period started with only water available, after which emotional behaviour was tested. Alcohol withdrawal did not induce major effects on anxiety or depression related behaviour in this study. Mitragynine (0, 5, 10 mg/kg; i.p.) did not affect anxiety-related behaviour or general locomotor activity in the open field test (Fig. 5E) or the elevated plus maze test (Fig. 5F). However, it had a significant dose-dependent antidepressant effect in the novelty suppressed feeding test (Fig. 5G) and the forced swim test (Fig. 5H). These data suggest that mitragynine may have beneficial effects on emotion during alcohol withdrawal, with antidepressant effects, but not anxiolytic effects.

### Mitragynine is self-administered and reduces alcohol consumption

A great problem for pharmacotherapies of addiction is their low compliance (McPheeters, et al., 2023). This may result from a pharmaco-development strategy in which all available treatments had been only experimenter administered and, in that way, yielded their therapeutic effects. In the human reality of an AUD, however, it is important to generate sufficient motivation for a therapeutic self-administration of the pharmacotherapy, of course without becoming addictive on its own terms. It is, therefore, postulated that any pharmacotherapy for AUD to become a candidate for further evaluation needs to be a.) self-administered (in a non-escalating way) and b.) this self-administration has to lead to a reduction in main AUD-related behaviours. Here we tested whether mitragynine would fulfil these criteria. We trained male mice to establish alcohol drinking in a social environment of the IntelliCage (n=12/cage). Thereby, one corner of the maze allowed nose-poke responding for alcohol with an FR1 plan, and one corner responding for tap water. After four weeks, a stable 16 vol.% alcohol consumption was established and then maintained for a further three weeks. Mitragynine (p.o.) self-administration was tested first by replacing the water bottle with mitragynine solution for one week (60 mg/l), with permanent free access for 6 days. This resulted in animals not consuming mitragynine, but maintaining their alcohol consumption (Fig. 5I). After mitragynine was removed again, animals were maintained with the alcohol/water drinking schedule for eight more weeks. Then a second mitragynine training was initiated. For one week, animals received for 6h/day only mitragynine solution in both accessible bottles. For the rest of the day (18 h), the alcohol/water schedule was maintained. In a subsequent 15-day test period, mitragynine remained available in one bottle and alcohol in the other for the whole time. We observed that animals now not only consumed a profound amount of mitragynine in a stable, non-escalating way over the 15 days, but at the same time significantly reduced their alcohol consumption (Fig. 5J). These findings may suggest that after some forced exposure training, animals not only learn to voluntarily self-administer mitragynine orally in a stable and non-escalating way, but at the same time reduce their alcohol consumption.

### Mitragynine brain binding

Mitragynine, once taken up into the blood, crosses the blood-brain barrier (Yusof et al., 2019). Although some binding partners of mitragynine in the brain are known, not all are fully acknowledged (Takayama et al., 2002; Matsumoto et al., 2004; Yusoff et al., 2025) and functionally validated yet (Yusoff et al., 2017, 2018; Gutridge et al., 2020, Japarin et al., 2023). In order to better understand the action in the brain, we traced mitragynine binding in the brain of mice. For this purpose, we developed and synthetized several fluorophore-marked mitragynine derivatives that showed an enrichment in slice preparations. The derivative displaying the most selective staining pattern, Mitragynine-Cy5, was then chosen for a more detailed analysis of mitragynine-associated fluorescence (Fig. 6A). Following ex vivo incubation of coronal brain sections, fluorescence was observed throughout the tissue, with particularly prominent signals in regions corresponding to major white matter tracts (Fig. 6B-C). To investigate this distribution in more detail, we examined hippocampal sections from two mice. In both animals, distinct Mitragynine-Cy5-positive structures were detected within the hippocampal tissue, while sections incubated with unconjugated Cy5 showed no comparable prominent focal staining in the selected regions (Fig. 6D-F).

**Figure 6.**
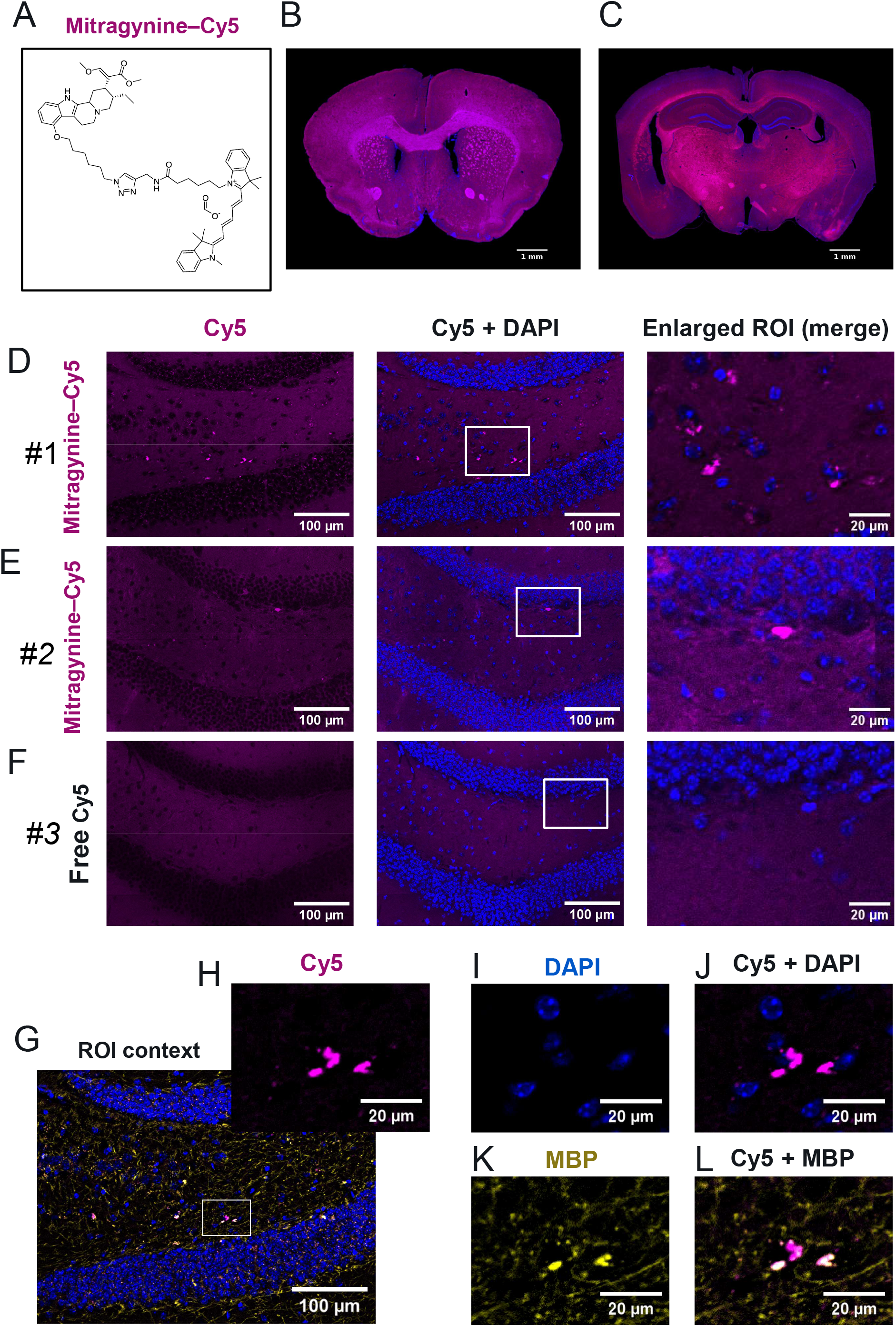
Ex-vivo fluorescence distribution of mitragynine in mouse brain sections and its spatial relationship with MBP immunoreactivity and DAPI-labelled nuclei. **A.** Chemical structure of the Cy5-conjugated mitragynine derivative Mitragynine-Cy5 (FGH484_1; molecular weight: 1075.409 g/mol). **B.-C.** Representative fluorescence images of coronal mouse brain sections incubated ex-vivo with FGH484_1 (10 µM), showing prominent Cy5 fluorescence in regions corresponding to major white matter tracts. **D.-F.** Representative hippocampal sections from two male C57BL/6J mice incubated with FGH484_1 (10 µM) or equimolar free Cy5 (10 µM). Each row shows Cy5 fluorescence (magenta), merged Cy5 and DAPI fluorescence (blue), and an enlarged view of the indicated region of interest (ROI), respectively. Discrete Cy5-positive structures are visible in FGH484_1-incubated sections, whereas the free-Cy5 control shows no comparable prominent focal fluorescence in the selected field. **G.-L.** Spatial distribution of FGH484_1-associated fluorescence relative to myelin basic protein (MBP) and DAPI-labelled nuclei in a hippocampal section. **G.** Overview showing Cy5 (magenta), MBP (yellow), and DAPI (blue), with the boxed region enlarged in H-L. **H.-J.** Cy5 fluorescence, DAPI-labelled nuclei, and the corresponding merged image, showing predominantly extranuclear Cy5-positive structures. **K.-L.** MBP immunoreactivity and the corresponding Cy5–MBP merged image, showing partial spatial overlap between Cy5-positive and MBP-positive structures. High-magnification images represent individual optical planes without Z-projection. Local background subtraction was applied to Cy5 and MBP channels for display in H-L. Scale bars: 1 mm (B, C), 100 µm (D-F, overview and merged images; G), and 20 µm (D-F, enlarged images; H-L).

We next examined whether these fluorescent structures were associated with myelin or cellular nuclei. For this purpose, sections were immunostained for myelin basic protein (MBP) and counterstained with DAPI (Fig. 6G-L). The analysis revealed that some Cy5-positive structures overlapped with MBP immunoreactivity, although a substantial proportion of the fluorescence was also detected outside MBP-positive regions. In one representative hippocampal optical plane, 141 of 584 Mitragynine-Cy5-positive structures (24.1%) fulfilled the criteria for spatial association with MBP. Pearson’s correlation coefficient between Mitragynine-Cy5 and MBP fluorescence intensities was 0.577 and decreased to 0.467 after background correction. In contrast, only 46 of 584 Mitragynine-Cy5-positive structures (7.9%) showed substantial overlap with DAPI-labelled nuclear regions, indicating that most detected structures were located predominantly outside the nuclei. The degree of overlap varied between the examined fields and optical planes.

These observations suggest that mitragynine-associated fluorescence is not restricted to regions containing previously described neuronal receptor targets, but is also found in white matter-rich structures. The partial spatial association with MBP suggests a possible interaction with components of myelinated tissue.

### Mitragynine attenuates tinnitus-related behaviour in Mongolian gerbils

The treatment of the animals was performed under two application regimes, first the acute treatment immediately after a monaural acoustic trauma with mitragynine (0, 5, 10 mg/kg) given by daily s.c. injections over three weeks. Second, the treatment started two weeks after the acoustic trauma at an early chronic state with corresponding groups and treatment duration (Ahlfs et al., 2012). In all cases, auditory brainstem response (ABR) hearing thresholds of the control and trauma ear as well as GPIAS behavioural tinnitus test data were acquired weekly. Thereafter, cochlear synapse count was obtained after sacrificing the animals at the end of treatment. In the acutely treated animals, mixed repeated-measures ANOVAs (factors: group and stimulation frequency, repetition: treatment weeks) of both ears revealed no significant interaction of all factors (trauma ear: Fig. 7A, 3 left panels; Suppl. Tab. 1). Nevertheless, group differences in hearing loss development over the three weeks of treatment could be found, with mitragynine treatment resulting in significantly worse hearing thresholds in trauma ears compared to the vehicle control (Fig. 7A, right panel). Corresponding GPIAS behavioural tinnitus tests did not show any significant changes in affected tinnitus animals over time in the vehicle group but a significant reduction in the number of animals showing tinnitus-related behaviour in both treatment groups after three weeks of mitragynine treatment (Bonferroni-corrected chi²-tests vs. corresponding vehicle group, p<0.05; Fig 7B). At inner hair cells (IHC), four of those animals did show a tinnitus-related behaviour at the first week of treatment and were, therefore, classified as tinnitus (T) animals, the other seven were classified as no-tinnitus (NT) animals. A two-way ANOVA (factors tinnitus status and cochlear frequency location) revealed a significant loss of IHC ribbon synapses only in the T animals, but not in the NT animals, after the trauma (Fig. 7C). This may suggest that, even though acute treatment with mitragynine over three weeks did not help in hearing loss recovery after the acoustic trauma, the animals were relieved of their tinnitus perception without any sign of synaptic protection or recovery.

**Figure 7.**
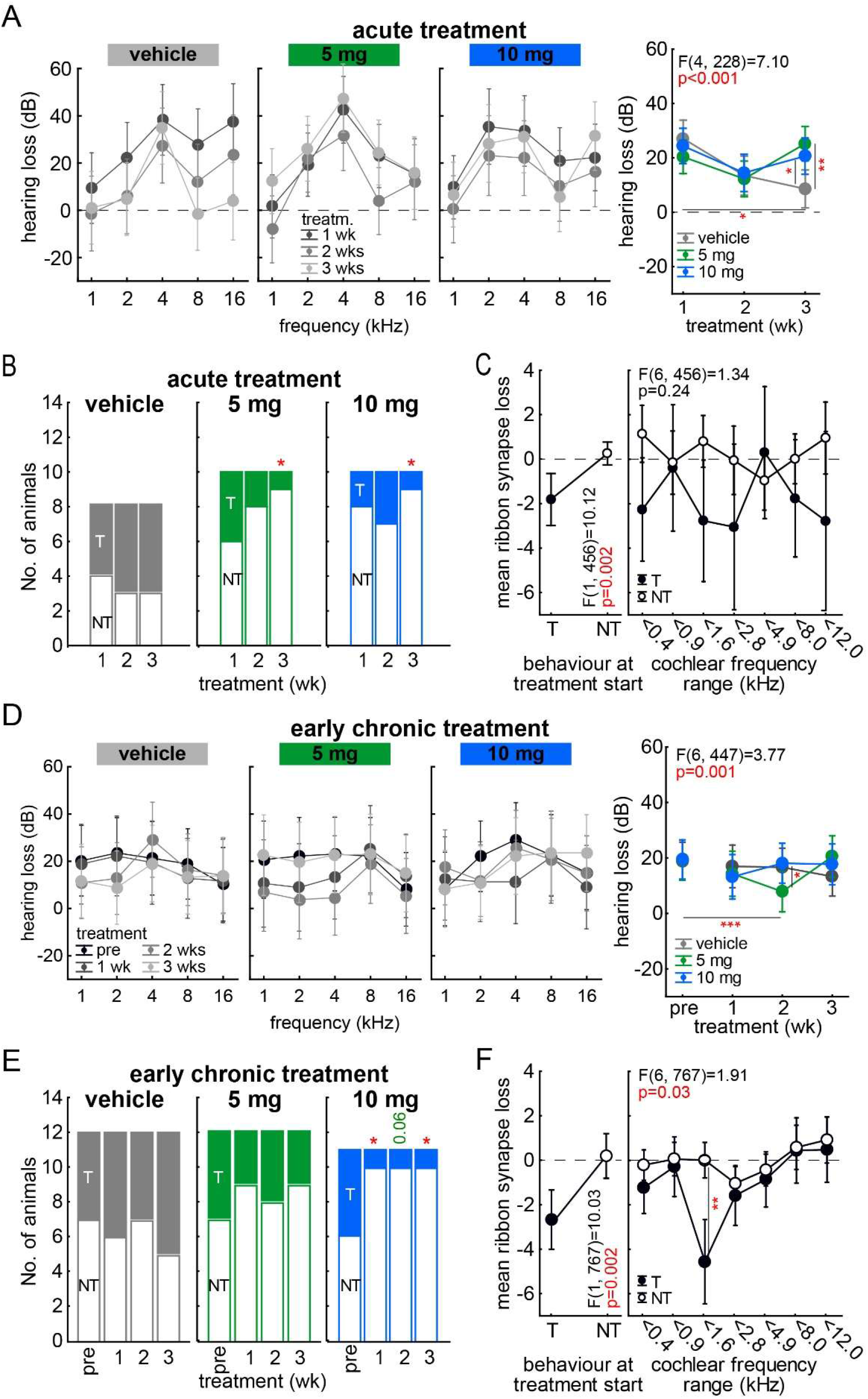
Mitragynine attenuates tinnitus after acute and chronic treatment in Mongolian gerbils. Effects of mitragynine on auditory brainstem response (ABR), gap prepulse inhibition of the ASR (GPIAS) and ribbon synapses of the inner hair cells (IHC) in acute (n=8-10/group) and early chronic (n=11-12/group) treatment regime. **A** Hearing loss (dB) in trauma ear over the three weeks of treatment in the three acutely treated animal groups (ANOVA; complete interaction: F_16,228_=1.23, p=0.24; interaction group and repetition: F_4,228_=7.10, p<0.001). Asterisks indicate significant Tukey post-hoc tests (*p<0.05, **p<0.01; for more statistical details: see Suppl. Tab. 1). **B** Number of animals with (T) or without (NT) tinnitus related behaviour over the acute treatment phase in the three treatment groups. Asterisks (*p<0.05) indicate Bonferroni-corrected chi²-tests vs. vehicle group. **C** Mean IHC ribbon synapse loss in acutely treated mitragynine animals in relation to the behaviour at treatment start (two-way ANOVA; behaviour: F_1,456_=10.12, p=0.002, cochlear frequency: F_6,456_=0.24, p=0.96; interaction: F_6,456_=1.34, p=0.24). **D** Hearing loss (dB) in trauma ear over the three weeks of treatment in the three early chronic treated animal groups (mixed repeated measurement ANOVA; complete interaction: F_24,447_=1.00, p=0.46; interaction group and repetition: F_6,447_=3.77, p=0.001). Asterisks indicate significant Tukey post-hoc tests (*p<0.05, *** p<0.001; for more statistical details: see Suppl. Tab. 2). **E** Number of T and NT animals before and over the early chronic treatment phase in the three treatment groups. Asterisks (*p<0.05) indicate Bonferroni-corrected chi²-tests vs. vehicle group. **F** Mean IHC ribbon synapse loss in early chronic treated animals in relation to the behaviour at treatment start (two-way ANOVA; behaviour: F_1,767_=10.03, p=0.002; cochlear frequency: F_6,767_=4.91, p<0.001; interaction: F_6,767_=1.91, p=0.03).

In the clinically more relevant early chronic treatment approach, we did not observe a significant increase of hearing loss after the acute treatment, but a significant transient decrease of hearing loss in the 5 mg mitragynine group that did not stabilize till the end of the treatment (Fig. 7D Suppl. Tab. 2). In comparison to the acute treatment, we only found a significant reduction of tinnitus related behaviour in the 10 mg treatment group with an early beginning already after the first week of treatment and staying till the end of the observation period (Fig. 7E). A two-way ANOVA of the IHC ribbon synapse loss showed the same pattern of increased synapse loss across all cochlear frequency locations in T animals only (Fig. 7F). Additionally, we found an interaction of both factors, indicating a frequency specific synaptopathy in T animals. Altogether, independent of any hearing loss changes, the higher dose of mitragynine was able to relieve the animals of tinnitus in an early chronic treatment approach from the start of the treatment on. However, lost synapses did not recover after the treatment.

## Discussion

Here we describe an inhibitory effect of mitragynine on various behaviours that define an AUD in specific scenarios that are characteristic of the disorder (Müller et al., 2025). We found that mitragynine inhibits well-established ongoing alcohol consumption in a social setting when alcohol is permanently available. Mitragynine also inhibits alcohol consumption in a non-social, isolated setting, though to a lesser degree. While mitragynine did not work for prophylaxis during abstinence in a social or non-social setting, it markedly attenuated the relapse to binge-drinking with alcohol availability restricted to a limited time. It also had a small beneficial effect when relapse had already started after a period of abstinence. Furthermore, mitragynine reduced alcohol seeking behaviour during its establishment as well as during retrieval from memory. It was also effective in attenuating a re-establishment after extinction. Since the aversive state during alcohol withdrawal is a key driver for relapse to alcohol drinking after abstinence, we investigated how mitragynine affects this situation. We found significant antidepressant effects of mitragynine, but no anxiolytic action, during alcohol withdrawal. A prevailing problem with AUD pharmacotherapies is the low compliance. Here we established a test paradigm that can monitor this and found that mice would not spontaneously take the treatment. After a short training, however, alcohol drinking mice learned to orally self-administer mitragynine at a constant rate. This went along with a strong reduction in voluntary alcohol drinking. To better understand the way mitragynine acts in the brain, we fluorescence-labelled mitragynine and tested its binding in brain slices. We did not find strong indication for binding at neurons where neurotransmitter receptors are localized, but strong enrichment in the white matter at myelin-containing fibres.

Here we report that mitragynine inhibits alcohol drinking behaviour in several distinct scenarios. Alcohol drinking is certainly at the centre of AUD, but occurs in quite different circumstances and with distinct motivations and goals (Müller et al., 2021, 2023, 2025). Its inhibition is, therefore, a crucial component of an AUD pharmacotherapy (Falk et al., 2019). We found an inhibition of ongoing drinking behaviour in a social context (social drinking) and in an isolated context (closer to depressed drinking) when alcohol was easily available. Inhibitory effects of Kratom and mitragynine on isolated drinking-in-the-dark were reported previously in male mice (Gutridge et al., 2020), and the use of Kratom with the intention to reduce alcohol consumption was reported in single cases in humans (Havemann-Reinecke, 2011; Singh et al., 2014, 2026; Coe et al., 2019). No inhibitory effect was observed on timely restricted drinking in the active phase, which closely resembles binge drinking in humans. Importantly, we found that alcohol-experienced constantly drinking animals can learn to orally self-administer mitragynine after a short, forced-exposure training. This self-administration went along with a subsequently reduced alcohol drinking in a social context. At the present stage, it is unclear whether this is an inhibitory effect of the mitragynine, or a drug-replacement effect, as it might work for morphine in animals (Japarin et al., 2021) and humans (Boyer et al., 2008; Grundmann et al., 2022). While the experimenter-administered mitragynine took place in a non-reinforcing dose (Yusoff et al., 2015, 2018; Japarin et al., 2023), the self-administered amount may have been higher at certain time points, which can have rewarding effects by itself (Yusoff et al., 2015). It should also be noted that a specific training with a forced exposure was required to initiate subsequent mitragynine self-administration. Nevertheless, no escalation in mitragynine oral self-administration was observed, which confirms the non-addictive property results from i.v. self-administration (Harun et al., 2015, 2020), intracranial self-stimulation (Behnood-Rod et al., 2020), and neurochemistry (Yusoff et al., 2022). To our knowledge, this is the first demonstration of a potential self-medication reducing alcohol consumption in rodents. It may open up a new pathway of treatment research in that it monitors compliance with an available pharmacotherapy and active reduction of a key symptom of the disorder. Evidence for zoopharmacognosis in animals and self-medication of physical impairments already exists, e.g., in monkeys (Page et al., 1992; Glander, 1994) and other species (Rodriguez and Wrangham, 1993; Huffman et al., 2003). This should be first evidence for self-medication of a mental disorder symptom.

As there are many distinct drinking scenarios in the temporal occurrence of AUD defining drinking occasions (Müller et al., 2025), we have tested several of them. Mitragynine did not work as a prophylactic treatment when given during withdrawal with the intention to attenuate subsequent relapse drinking in a social context. It rather enhanced the relapse when tested in isolated drinking, but could attenuate the relapse to binge-drinking. Once relapse has commenced, a pharmacotherapy is meant to reduce the drinking amount during the relapse. In this scenario, we found no effectiveness for mitragynine. Mitragynine had no major effect on water consumption, suggesting that it may not interfere with general fluid regulation of the organism.

Altogether, these observations would make the use of mitragynine as a potential therapeutic option phase-sensitive (Müller et al., 2025), i.e. it may be only advocated for certain drinking phases, but not for unambiguous administration.

A major component of AUD is alcohol seeking behaviour, which is driven by internal and/or external cues, or is spontaneously occurring. It brings the affected individual close to an alcohol source and allows for the initiation of drinking behaviour (Bach et al., 2025, 2019; Hoffmann et al., 2023). Here we report that mitragynine can inhibit the establishment as well as the retrieval of established alcohol seeking behaviour in mice. This effect was dose-dependent and manifested over the entire learning curve. We also found that it can reduce the re-establishment of already extinguished alcohol seeking behaviour. This profile would add to a therapeutic effect of mitragynine in AUD as drug memories are inhibited in their build-up as well as their retrieval (Robbins et al., 2008; Müller and Schumann, 2011).

A highly aversive state is experienced by AUD patients during withdrawal. Also, this is an AUD criterion (Koob and Mason, 2016). It was argued that this negative emotional state, characterized by anxiety, anhedonia, and depression, is a major driver for relapse to alcohol consumption (Heilig et al., 2010; Koob and Volkow, 2016). Here we found that experimenter-administered mitragynine attenuates depression-like behaviour in the novelty supressed feeding test and the forced swim test during withdrawal from chronic alcohol drinking. However, it had no effects on anxiety-related behaviour in the open field or elevated plus maze test. It should be noted that withdrawal effects in the study appeared as rather mild. Nevertheless, mitragynine can make the withdrawal phase less aversive and, by that way, attenuate relapse likelihood (Day and Day, 2022). Mitragynine was previously reported to also attenuate withdrawal effects of morphine in rats (Hassan et al., 2020), and Kratom may have a general antidepressant profile in humans (Johnson et al., 2020; Timmons et al., 2026). However, long-term mitragynine application may also yield some withdrawal effects on its own (Hassan et al., 2021, Yunusa et al., 2024).

The effects of mitragynine on the sedative effects of alcohol were ambiguous in this study. After acute treatment, it delayed sedation, but enhanced its duration. Repeated high-dose alcohol exposure leads to tolerance development: the latency for sedation increases, and duration declines. Together with this tolerance development, mitragynine loses its efficacy to delay the onset, but still elongates the time of sedation. While the sedating properties of alcohol are not a direct AUD criterion, they are, nevertheless, predictive of the risk of establishing an AUD (Spanagel, 2009). Enhanced sedation time, however, should reduce this in line with a therapeutic application.

Mitragynine is the major psychoactive compound of the plant preparation Kratom. Various studies investigated the effects of Kratom extracts on AUD-related behaviours in rodent models and found that it can reduce alcohol seeking behaviour (Vijeepallam et al., 2019), diminish alcohol intake (Gutridge et al., 2020), and attenuate alcohol withdrawal symptoms (Kumarnsit et al., 2007; Cheaha et al., 2015). However, Kratom extracts were of variable, often unknown composition with other potent psychoactive components, such as 7-OH-mitragynine (Gutridge et al., 2020; Prevete et al., 2022).

For a new therapy of AUD, a potential neuropharmacological mechanism may be desirable. In order to identify this, we determined brain binding of fluorescence-labelled mitragynine. This was mainly with the intention, to possibly confirm and extend its previously suggested partial agonist action at µ-opioid receptors in the brain (Bhowmik et al., 2021; Kruegel et al., 2016). Much to our surprise, we found the majority of binding not at neuronal membranes where µ-opioid receptors are expected to be localized, but in large parts of the white matter. This may suggest a previously unrecognized mechanism of mitragynine in its anti-AUD action, that, still has to be further explored (Haile et al., 2026).

A motivation to test for mitragynine as a therapeutic substance was a single-case patient report, where a patient claimed self-medication and therapeutic success to treat not only a confirmed AUD, but also a tinnitus (Müller et al., 2023). This is certainly unusual, as AUD and tinnitus are not typically occurring together as comorbidities. However, they may both be both driven by joint environmental factors, such as stress and development of a depression (Spanagel et al., 2009; Müller et al., 2023; Patil et al., 2023). This study also investigated the effects of mitragynine on tinnitus in Mongolian gerbils. Neither in the ABR hearing thresholds nor in the extent of the acoustic trauma-induced synaptopathy at the level of the IHC, was any benefit found after mitragynine treatment. Other studies on hearing loss (Nieratschker et al., 2024) and tinnitus (Tziridis et al., 2025) found improvements after acute substance application. In these cases, both, hearing thresholds and IHC synaptic counts, could be protected from cochlear damage. While the protective effect of acute treatment on tinnitus development (Tziridis et al., 2025) probably resulted from protection from trauma-induced synaptopathy, the effect of mitragynine on tinnitus rather seems to be a centrally mediated. Clinically even more relevant is our observation that, in contrast to other substances tested, mitragynine was also effective in relieving tinnitus in the early chronic condition. Interestingly, in the acute treatment, both substance doses were able to decrease tinnitus-related behaviour after three weeks of application, while in the early chronic treatment, only the 10 mg mitragynine group showed a significant reduction of tinnitus-related behaviour from the beginning of the treatment. In both conditions, the question whether the observed reduction in tinnitus-related behaviour is a consequence of a reduction in tinnitus loudness, or a general improvement of mood, or other central changes, cannot be answered based on our current results and would need further research.

There are limitations to this work that may still restrict interpretation of the results. First, AUD is a mental disorder that seldom occurs alone. In fact, it is often established in individuals with preceding or co-developing other mental disorders, partly in order to use alcohol as self-medication (Müller et al., 2023). This therapeutic efficacy may preferentially be needed in comorbid organisms (e.g. with depression and anxiety; Müller et al., 2017; Müller and Kornhuber, 2017). This was not tested here and remains to be determined in the future. Second, a self-medication with mitragynine was only shown in relation to alcohol drinking behaviour, but not with other AUD-related behaviours. For alcohol drinking, only one specific (mitragynine-sensitive) scenario - alcohol drinking in a social context – was tested. Although this effect would, in our eyes, already justify the pursuit of mitragynine as an AUD therapy, it would be important to know how mitragynine self-medication compliance holds up for other AUD behaviours. Third, also a self-mediation with mitragynine that leads to a reduction of the tinnitus symptoms may be desirable, before advocating the compound, in particular in the light of the fact that there is currently no pharmacotreatment available for tinnitus therapy. Fourth, this study investigated only males. Although AUD predominantly affects males, there is a growing number of females suffering from this disorder. Recent findings focusing on females, however, also showed promising therapeutic effects of mitragynine on operant alcohol self-administration in female rats (Haile et al., 2026). Thus, present results need to be replicated to the full extent in females.

Altogether, the present results suggest that mitragynine is able to reduce several key symptoms of AUD in rodent models in which the compound is orally self-administered, showing a good compliance. Furthermore, it may reduce behavioural symptoms of tinnitus.

## Methods

### Animal studies

#### Animals

All experiments were carried out in accordance with the National Institutes of Health guidelines for the humane treatment of animals and the European Communities Council Directive (86/609/EEC) and approved by the local governmental commission for animal health (German state administration Bavaria/ Regierung von Unterfranken). Male C57/Bl6J mice, 8-10 weeks old, weighing 20-28 g at the beginning of the experiment, were used. Animals were housed in groups in standard macrolone cages (Type III, Macrolone®), or individually housed in these cages in drinking studies. They were provided with food and water ad libitum, with nesting material as cage enrichment, and kept on a 12:12-hour light: dark cycle. Behavioral tests were performed during the light cycle between 09:00 and 16:00 h. Room temperature was maintained between 19°C and 22°C at a humidity of 55% (±10%).

Male Mongolian gerbils (*Meriones unguiculatus*), purchased from Janvier (Le Genest-Saint-Isle, France) were housed in standard Type IV macrolone cages in a ventilated cabinet (Bio A.S. Vent Light, Zoonlab, Emmendingen, Germany) in groups of 3-4 animals with free access to water and food at a room temperature of 20 to 25°C under a 12:12-hour dark/light cycle. At the beginning of the experiments, the animals were 12 weeks old and had been at least two weeks in the animal facility for habituation.

### Drug preparation

Mitragynine extraction, isolation and verification from the fresh leaves of *Mitragyna speciosa* were conducted at the Centre for Drug Research (CDR), Universiti Sains Malaysia (USM), using a previously described method (Utar et al., 2011). Mitragynine purity was verified by utilizing both high-performance liquid chromatography (HPLC) and proton nuclear magnetic resonance (1H-NMR) (400 MHz) analysis. The process yielded mitragynine with a 98% purity (Yunusa et al., 2024; Suppl. material). Mitragynine was dissolved in vehicle (20% (v/v) Tween 80 diluted with physiological saline; 0.9 % NaCl; Sigma Aldrich, UK).

For the mitragynine self-medication study, a water-soluble preparation was used. Mitragynine hydrochloride was prepared from mitragynine base using an in-house conversion procedure to obtain a water-soluble salt suitable for experimental administration. The purity of the resulting salt was determined using high-performance liquid chromatography (HPLC) and was found to be greater than 99%. The characterized mitragynine hydrochloride was subsequently used in the animal self-administration study.

### Social alcohol drinking with intermittent access

In this experiment, we tested the efficacy of mitragynine to reduce alcohol consumption in a social setting, a.) during ongoing intermittent access drinking, and b.) as a relapse prevention (König et al., 2020). Fourteen male C57Bl/6J mice were all transferred to an IntelliCage (TSE systems, Bad Homburg) and maintained there for the rest of the experiment. They were provided with food and water ad libitum, with red transparent triangular shelters and nesting material as cage enrichment. The IntelliCage contains four corners, with operant boxes for nose pokes and a fluid dispenser in each. During the drinking phase, all corners contained alcohol for 2h/day (9-11 am) or water (22h/day), both with a fixed ratio 1 (FR1) access schedule. Individual responses of the animals were measured by a chip contact at the nose poke and liking units of the corners (Pagano et al., 2023; Salamian et al., 2026).

After a seven-day acclimatization period to the IntelliCage, alcohol consumption was established in all mice. Animals received alcohol at increasing concentrations of 3, 6, 12, and 20 vol.% in the corners of the maze. Mice were exposed to each concentration of alcohol for 4 days. Thereafter, alcohol concentration was maintained at 20 vol.% and animals were allowed to drink for 4 weeks. In order to measure the alcohol deprivation effect, alcohol was removed for 3 weeks (all bottles containing tap water) before it was reintroduced with a concentration of 20 vol.% for 4 days. This procedure was repeated once more. Bottles were changed and weighed daily. The consumed amount of alcohol and water relative to body weight was measured (Pagano et al., 2023; Salamian et al., 2026).

After 20 vol.% alcohol drinking was established, animals were divided into 2 groups (n=7/group). Each group received four treatment episodes: i.) & ii.) during ongoing drinking, iii.) and iv.) during withdrawal/before reinstatement and expression of an alcohol deprivation effect (ADE) (König et al., 2020). A treatment episode consisted of three days with one i.p. injection/day (v_inj_=10 ml/kg) of either mitragynine (5 mg/kg) or vehicle, administered 1 h before alcohol was introduced every day. The mitragynine dose was chosen according to a previous report where this dose regimen showed no reinforcing effects on its own (Yusoff et al., 2015), but reduced reinforcing effects of morphine in rodents (Japarin et al., 2021). The single group schedule of treatment episodes was: group 1: mitragynine-vehicle-mitragynine-vehicle and group 2: vehicle-mitragynine-vehicle-mitragynine. We did not observe statistical differences between groups at each treatment episode. Therefore, groups were collapsed for analysis at respective treatments and time points.

### Isolated alcohol drinking with unlimited access

In this experiment, we tested the efficacy of mitragynine to reduce alcohol consumption in single-housed isolated setting, a.) during ongoing unlimited access drinking, b.) as a relapse prevention, and c.) as a relapse-limiting treatment (König et al., 2020). 33 male C57Bl/6J mice, 8 weeks old, were all single-housed in standard cages and maintained there for the rest of the experiment. They were provided with food and water ad libitum, with nesting material as cage enrichment. After a 14-day acclimatization period with two bottles available (tap water), alcohol consumption was established in all mice. Animals received alcohol at increasing concentrations of 2, 4, 8, 12, and 16 vol.% in one of the bottles. The other bottle always contained water. Mice were exposed to each concentration of alcohol for 4 days. Thereafter, alcohol concentration was maintained at 16 vol.% and animals were allowed to drink for 3 weeks. In order to measure the alcohol deprivation effect, alcohol was removed for 3 weeks (all bottles containing water) before it was reintroduced with a concentration of 16 vol.% for 4 days. This procedure was repeated one more time. Bottles were changed and weighed daily. The consumed amount of alcohol and water relative to body weight and the preference for alcohol vs. water were measured (Easton et al. 2013; Müller et al. 2017).

After 16 vol.% alcohol drinking was established, animals were divided into 3 groups (n=11/group). Each group received three treatment episodes: i.) during ongoing drinking, ii.) during withdrawal/before reinstatement of ADE, and iii.) during reinstatement of ADE (König et al., 2020). A treatment episode consisted of three days with one i.p. injection/day (v_inj_=10 ml/kg) of either mitragynine (5 mg/kg) or vehicle, administered at 6 pm on the respective days. The mitragynine dose was chosen according to a previous. The single group schedule of treatment episodes was: group 1: mitragynine-vehicle-vehicle, group 2: vehicle-mitragynine-vehicle, and group 3: vehicle-vehicle-mitragynine. We did not observe statistical differences between groups at each treatment episode. Therefore, groups were collapsed for analysis at respective treatments and time points.

### Alcohol binge-drinking with limited access

Alcohol binge drinking and mitragynine effects were tested in mice in three different drinking scenarios of a drinking-in-the-dark (DID) paradigm: a.) during ongoing unlimited access drinking, b.) as a relapse prevention, and c.) as a relapse limiting treatment (König et al., 2020). 33 male C57Bl/6J mice, 8 weeks old, were all single-housed in standard cages and maintained there for the rest of the experiment. They were provided with food and water ad libitum, with nesting material as cage enrichment, and kept on a reversed 12:12-hour light: dark cycle (lights on at 7.00 pm). After a seven-day acclimatization period with two bottles available (tap water), alcohol consumption was established in all mice for 3 weeks. Animals received alcohol at 20 vol.% in one of the bottles only during the dark phase (7 am - 7 pm). In order to measure the alcohol deprivation effect, alcohol was removed for two weeks (all bottles containing water) before it was reintroduced for eight days. This procedure was repeated once more. Bottles were changed and weighed daily. The consumed amount of alcohol and water relative to body weight and the preference for alcohol vs. water were measured (Stürzemberger et al., 2026).

After 20 vol.% alcohol DID was established, animals were divided into 3 groups (n=11/group). Each group received three treatment episodes: i.) during ongoing drinking, ii.) during withdrawal/before reinstatement of ADE, and iii.) during reinstatement of ADE (König et al., 2020). A treatment episode consisted of three days with one i.p. injection/day (v_inj_=10 ml/kg) of either mitragynine (5 mg/kg) or vehicle, administered at 6 am on the respective days. The mitragynine dose was chosen according to the previous experiments. The single group schedule of treatment episodes was: group 1: mitragynine-vehicle-vehicle, group 2: vehicle-mitragynine-vehicle, and group 3: vehicle-vehicle-mitragynine. We did not observe statistical differences between groups at each treatment episode. Therefore, groups were collapsed for analysis at respective treatments and time points.

### Conditioned place preference

The establishment and expression of conditioned place preference (CPP) was tested in naïve male C57Bl/6J mice, 8 weeks old at the beginning of the experiment. The TSE Place Preference test boxes (Bad Homburg, Germany) were made of non-transparent PVC with standard inside dimensions of 40cm (L) x 15cm (W) x 20cm (H). The apparatus was divided into 3 fully automated compartments; the outer chambers measured 17cm in length, and the centre chamber 6 cm. The floor of the left chamber (compartment A) was covered with a smooth black rubber mat. The floor of the right chamber (compartment B) was covered with a patterned black rubber mat. The centre chamber was not covered and was coloured white (compartment C). Activity was recorded in each compartment using high-resolution infrared sensors. The system automatically recorded the number of entries made, the sojourn time, and distance moved in each compartment for each trial. An unbiased design was used, i.e. half the mice were conditioned to their preferred compartment, and half to their non-preferred compartment (Müller et al., 2017; Kalinichenko et al., 2021). Animals were injected (i.p.) immediately before each trial with either saline or alcohol (2 g/kg). Mice were instantly transferred to the testing suite and placed into the CPP boxes, signifying the beginning of the trial period. The experiment involved four phases: habituation trial (one session), baseline testing (Bl), conditioning trials (14 sessions), and preference tests (3 sessions, T1-T3). Trials were performed once daily.

#### Habituation

The habituation session was intended to acclimatise mice to the test procedure and apparatus prior to commencing the experiment. Mice were injected with saline and introduced into the centre compartment with free access to all three compartments for 20 minutes. *Baseline-test:* The pre-test was designed to establish a baseline level of preference for each individual animal. Mice were injected with saline and introduced into the centre compartment with free access to all three compartments for 20 minutes.

#### Conditioning trials

Conditioning trials were performed in pairs, odd-numbered pairings were conditioned with alcohol, and even-numbered pairings were conditioned with saline, this was balanced across groups. All animals received 7 pairings with saline and 7 pairings with alcohol. Mice were injected with either saline or an ethanol solution and introduced into one of two compartments, with restricted access, for 5 minutes.

#### Preference tests

To monitor the time course of CPP establishment, preference tests were systematically performed after one, three and seven conditioning trials. Before each test, mice were injected with saline and introduced into the centre compartment with free access to all three compartments for 20 minutes (Müller et al., 2017; Kalinichenko et al., 2021).

#### Establishment

To test mitragynine effects on alcohol CPP establishment, three groups of naïve animals (n=11/group) received an i.p. injection with either mitragynine (5 or 10 mg/kg; i.p.) or vehicle, 30 min before each conditioning trial.

#### Expression

To test mitragynine effects on CPP expression (retrieval), three groups of naïve animals (n=11-12/group) received an i.p. injection with either mitragynine (5 or 10 mg/kg; i.p.) or vehicle, 30 min before each test trial.

#### Extinction

In order to test mitragynine effects on CPP extinction, two groups of naïve animals (n=14-15/group) received an alcohol CPP training and testing. Thereafter, they received an extinction training for 5 days, during which they were exposed to the unrestricted maze for 5 min. This was followed by two re-instatement training sessions with alcohol (2 mg/kg; i.p.) and saline, otherwise as above. In addition, they received an i.p. injection with either mitragynine (5 mg/kg; i.p.) or vehicle 30 min before each trial. One day later, they underwent a reinstatement test trial as above.

### Loss of righting reflex

#### Acute

Naïve male C57BL/6J mice, 8 weeks old at the beginning of the experiment were divided into three groups (n=12/group) and received a single i.p. treatment with mitragynine (0, 5, or 10 mg/kg). Thirty minutes later, they were injected with 3.5 g/kg (i.p.) alcohol to induce a loss of the righting reflex (LORR), and immediately placed in an empty cage. LORR was registered after the animal became ataxic and stopped moving for at least 30 seconds. Then the animal was placed on its back. Recovery from alcohol administration was considered when the animal was able to right itself three times within a minute. A 2h cut off was used. Time taken for the animal to lose its righting reflex, and time to recovery from alcohol’s effect were recorded (Zheng et al., 2016; Kalinichenko et al., 2021).

#### Tolerance

Naïve male C57BL/6J mice, 8 weeks old at the beginning of the experiment were divided into three groups (n=12/group) and were tested for a LORR as described above. Thereafter, they received a daily alcohol treatment (3.5 g/kg, i.p.) for seven days, in order to develop tolerance to the sedating effects. On day eight, they received a single i.p. treatment with mitragynine (0, 5, or 10 mg/kg). Thirty minutes later, they were injected with 3.5g/kg (i.p.) alcohol to re-test the LORR.

### Withdrawal-related emotional behaviour

Three groups of naïve male C57Bl/6J mice, 8 weeks old at the beginning of the experiment, were trained to drink 10 vol.% alcohol in a DID paradigm as described above. A control group always drank only water (n=10/group). After six weeks of drinking during which stable and comparable levels of consumption were established by all groups, animals underwent a two-week withdrawal, during which only water was available (Holleran and Winder, 2017). Thereafter, emotional behaviour was tested after treatment with mitragynine (0, 5, 10 mg/kg; i.p.) 30 min before each test. Mice were tested using a battery of behavioral tests in the following order: open field, elevated plus maze, novelty suppressed feeding, forced swim test, and sucrose preference tests. Mice were tested in a pseudorandom order and were moved to the behavioral suite adjacent to the housing room at least 1 hour before testing. Each test apparatus was cleaned with 70% ethanol between subjects to avoid any olfactory cues influencing behaviors. Mice were returned to their home cages at the end of each test and allowed to recover for at least 2 days before further testing. Behaviors for all tests were recorded for subsequent scoring (Müller et al., 2017; Kalinichenko et al., 2023).

#### Open Field

Each mouse was placed in a square white acrylic arena (50×50 cm), facing an outer wall, for 20 min and allowed to freely explore the arena. White light of 100 lx was evenly distributed across the arena during testing. Video recordings were taken and analyzed using Biobserve Viewer III (Biobserve GmbH, Germany). A virtual square of equal distance from the periphery (36×36 cm) was defined as the „central zone‟ in order to determine the number of entries, and time (s) spent in the central zone. Distance moved in the outer and central zones (cm), number of entries and time spent in the central zone were registered.

#### Elevated Plus Maze

The elevated plus maze was constructed from black opaque acrylic with white lining on the floor, each arm measuring 30×5 cm and the central platform 5×5 cm. One set of arms, opposing one another, was enclosed completely by a wall of opaque acrylic, 15 cm high, while the other set was open with a ledge of 0.5 cm on either side of the arms. The open arms were brightly lit (100 lx). The maze was elevated 50 cm from the ground on a transparent acrylic stand. Each mouse was placed on the central platform, facing towards a closed arm, and allowed to freely explore the maze for 5 min. Biobserve Viewer III tracking software (Biobserve GmbH, Germany) was used to record locomotor activity during the test (distance moved in the open and closed arms), and the number of entries into the closed and open arms and time spent in them. An arm entry was counted when two paws had entered an arm, and an arm exit was determined when two paws had left the arm.

#### Novelty suppressed feeding

Animals were deprived of food for 24 h before novelty-suppressed feeding test. After food deprivation, each mouse was put in the corner of a square white acrylic arena (50×50×50 cm), facing an outer wall. White light of 100 lx was evenly distributed across the arena during testing. A piece of food was placed in the centre of the arena. Video recordings were taken and analyzed using Biobserve Viewer III (Biobserve GmbH, Germany). The time (s) before a mouse began eating after the fasting period (24 h) and the distance moved before eating were registered. *Forced swim test:* For a short version of the forced-swim test, each mouse was placed into a glass transparent cylinder (17 cm diameter, 18 cm height) filled with water (12 cm, 25°C) for 6 min. Then an animal was returned to the home cage. The latency to first floating and total floating time were recorded manually.

#### Sucrose preference test

Animals were single-housed and had access to two bottles with water or a 0.5% sucrose solution for 2 days. The weight of animals was measured before and after the test, and the volume of water and sucrose solution was estimated daily. Sucrose preference in % of the drinking fluid was calculated (Müller et al., 2017; Kalinichenko et al., 2023).

### Mitragynine therapeutic self-administration

Male C57Bl/6J mice were trained to drink alcohol in the social environment of an IntelliCage (n=12/cage). Thereby, two corners of the IntelliCage were active with an FR1 plan. Each corner contained two bottles with water available all the time. After two weeks of adaptation to the IntelliCage, water in one of the bottles was replaced by alcohol at concentrations of 2, 4, 8, 12, and 16 vol.%. Each concertation was offered for four days, respectively. A stable 16 vol.% alcohol consumption was established and then maintained for a further two weeks (Pagano et al., 2023, Salamian et al., 2026). Mitragynine (p.o.) self-administration was tested first by replacing one water bottle with mitragynine solution (60 mg/l in tap water) for one week, with permanent free access for 7 days. The second water bottle as well as two bottles with alcohol were constantly available. This resulted in animals not consuming mitragynine, but maintaining their alcohol consumption. After mitragynine was removed again, animals were maintained with the alcohol/water drinking schedule for eight more weeks. During this time, each animal received two mitragynine injections (10 mg/kg, i.p.) spaced two weeks apart, to familiarize animals with the discriminative stimulus effects of the substance (Harun et al., 2015). Then a second mitragynine training was initiated. For one week, animals received for 6h/day only mitragynine solution placed in bottles previously used for alcohol in both accessible corners. For the rest of the day (18h), the alcohol/water schedule was maintained. In a subsequent 16-day test period, mitragynine remained available in one bottle and alcohol in the other for the whole time.

### Mitragynine brain distribution

#### Brain tissue preparation

Mice were anaesthetized with isoflurane and transcardially perfused with 0.9% NaCl followed by 4% paraformaldehyde (PFA). Brains from two mice (12–13 weeks old) were used for ex vivo fluorescence analysis. Following overnight post-fixation in 4% PFA at 4 °C, brains were cryoprotected in 30% sucrose in 0.1 M phosphate buffer. Coronal sections (40 µm) were stored in cryoprotectant at −20 °C until use.

#### Ex vivo labelling and immunofluorescence

Free-floating sections were washed in Tris-buffered saline (TBS) and blocked for at least 1 h at room temperature in TBS containing 10% normal donkey serum (Sigma-Aldrich, D9663) and 0.3% Triton X-100. Sections were incubated for 72 h at 4 °C with compound FGH484_1 (Mitragynine-Cy5) or unconjugated Cy5 (10 µM each) and rat anti-MBP antibody (Bio-Rad, MCA409S; 1:250). FGH484_1 was dissolved in DMSO before dilution in blocking solution. Following TBS washes, sections were incubated overnight at 4 °C with Cy3-conjugated donkey anti-rat IgG (Jackson ImmunoResearch, 712-165-153; 1:1,000), counterstained with DAPI (1:10,000; 15 min), and mounted using ProLong Gold Antifade Mountant.

#### Fluorescence microscopy

Images were acquired using a ZEISS Axio Imager. M2 microscope equipped with an ApoTome module and AxioCam MR R3 camera (ZEN blue software). Images were acquired at 10× and 20× magnification as 16-bit multichannel images. At 20× (Plan-Apochromat 20×/0.8), Z-stacks comprised ten planes at 1.5 µm intervals (0.3225 µm/pixel). Representative images show individual focal planes without Z-projection. Cy5, MBP (Cy3), and DAPI were pseudocoloured magenta, yellow, and blue, respectively.

#### Quantitative fluorescence intensity correlation and spatial overlap analysis

Cy5-MBP co-localization and Cy5-MBP/DAPI spatial overlap were analyzed in Fiji (ImageJ) using custom Groovy scripts. Original 16-bit images were analyzed within defined regions of interest (ROIs), with each Z-plane processed independently without projection. Pearson’s correlation coefficients for Cy5 and MBP were calculated from raw and background-corrected, non-thresholded intensities. Object-based spatial overlap was assessed separately. Local background and intensity variation (F; 1.4826 × local median absolute deviation, minimum 1) were estimated using 17 × 17-pixel median filtering after fourfold downsampling. Foreground detection required a background-corrected intensity ≥5F and a raw-to-background ratio ≥2. Eight-connected components smaller than four pixels were excluded. Cy5 components were classified as MBP-associated when ≥75% of their area overlapped the MBP mask and the mean MBP intensity met both thresholds. DAPI-labelled nuclei were segmented using Gaussian smoothing (σ = 1.2 pixels), background subtraction (σ = 24 pixels), Li thresholding, and morphological processing, excluding nuclear regions outside 20– 10,000 pixels. Cy5 components were considered nucleus-associated when ≥75% of their area overlapped the nuclear mask. Threshold sensitivity was assessed using 81 parameter combinations for Cy5-MBP and alternative Otsu-based nuclear masks for Cy5–DAPI. Results were interpreted descriptively without treating images, Z-planes, or fluorescent components as independent biological replicates (n=2).

### Tinnitus model in Mongolian gerbils

All animals were handled over the time course of one week before the beginning of the experiments and accustomed to the setup environments to minimize stress. Animals were separated into six groups with at least 8 animals/group. The groups consisted of 8 vehicle animals and two substance groups (5 mg: 10 animals; 10 mg: 10 animals) in the acute treatment approach and 12 vehicle animals and two substance groups (5 mg: 12 animals; 10 mg: 11 animals) in the early chronic experiments. The experimenters were blinded to the content of the medium applied. All methods used have been outlined elsewhere (Tziridis et al., 2021) and are therefore only briefly described below (Suppl. Fig. 1). Briefly, injections were applied s.c. by 15 daily applications over three weeks, either starting immediately after a mild monaural acoustic noise trauma or two weeks after that. Before any treatment, animals were behaviourally and electrophysiologically evaluated to obtain the baseline conditions for the tinnitus test and the auditory brainstem response (ABR) based hearing threshold measurements. Over the three weeks of application, the measurements were repeated once a week and in the case of the early chronic groups, once before treatment started.

After the end of the treatment, the animals were sacrificed, both cochleae were extracted, and IHC ribbon synapse numbers were evaluated immunohistologically.

#### Behavioral tinnitus assessment (by GPIAS)

The acoustic startle reflex (ASR) is a widely used behavioural approach that does not require any training before testing. It has been modified into gap prepulse inhibition of the ASR (GPAIS) to assess a possible tinnitus percept (Gerum et al., 2019; Schilling et al., 2017). In this study, animals were tested first, with a gap – no-gap paradigm to measure the prepulse inhibition (PPI) of the ASR amplitudes, one to two days before the acoustic trauma. These data were used as the healthy baseline condition for later calculation of the effect size of the change in PPI. For ASR measurement, animals were placed in a 15 cm long, 4.2 cm wide acrylic tube on a custom-made 3D-acceleration sensor platform in front of two speakers (one for the background stimulus, one for the startle pulse) inside a sound-attenuated, anechoic, dark chamber. For habituation, the animals were allowed 15 min of rest, followed by stimulation with five stimuli that were discarded from further analysis. The proper GPIAS stimuli consisted of band-pass filtered background noise (mean duration of 10 s; 60 dB SPL) with centre frequencies of 1, 2, 4, 8, and 16 kHz and half an octave bandwidth each. Each stimulus was presented 60 times, with (30 stimuli) or without (30 stimuli) a 20 ms gap, 100 ms before a startle stimulus (white noise burst, 20 ms duration, 2 ms sin²-ramp, 105 dB SPL) in a pseudorandomized manner. A complete behavioural measurement of 300 stimuli had a duration of roughly 45 min. The same measurements were repeated several times after the monaural acoustic trauma at the same time of the day.

#### Assessment of hearing thresholds (by ABR)

One day after each GPIAS measurement, the animals were anesthetized with 0.3 to 0.4 ml of a mixture of ketamine (75 mg/kg) and medetomidine (0.5 mg/kg) and placed on a remote-controlled heating pad inside a sound-attenuated, anechoic, dark chamber to obtain individual ABR-based audiograms (Schilling et al, 2019). The stimulation frequencies between 1 and 16 kHz in octave steps for stimulation intensities ranging from 0 to 90 dB SPL in 5 dB steps were presented free-field via a loudspeaker placed 3 cm in front of the pinna of the measured ear. For each ear, stimulus, and intensity, 300 repetitions of pairs of 6 ms long (each) phase-inverted double stimuli (100 ms intrastimulus interval; 500 ms interstimulus interval) were presented. The complete measurement of one ear took less than 30 min.

For ABR measurements, three silver electrodes were placed subcutaneously, one retroaural above the bulla of the tested ear (recording electrode), another one central between both ears (reference electrode) and a last one at the base of the tail (ground electrode). The signal was recorded and filtered (bandpass filter 400 to 2000 Hz) via a Neuroamp 401 amplifier (JHM, Mainaschaff, Germany). As for the GPIAS, the exact same measurements were repeated several times after the monaural acoustic trauma at the same time of the day.

#### Monaural acoustic trauma

A few days after the baseline measurements of GPIAS and ABR, the animals were again anesthetized with the same ketamine-medetomidine mixture and placed on a remote-controlled heating pad, 10 cm in front of a speaker aligned to their left ear inside a sound-attenuated, anechoic, dark chamber. The right ear (intra-individual control) was muted with protective ear foam (Ohropax®). A mild monaural acoustic trauma was induced in the left ear using a 2 kHz pure tone at 115 dB SPL lasting for 75 min (Tziridis et al., 2021). The animals were sleeping for approx. 120 min and were returned to their home cage after fully waking up.

The obtained data of the GPIAS and ABR measurements were evaluated objectively and automatically by custom-made Python programs (Schilling et al., 2019; Gerum et al, 2019). Tinnitus development was tested for each animal and frequency individually by t-tests of the log-normalized PPI: The log-normalization of the response amplitudes of gap (Agap) and no-gap (Ano-gap) by log(Agap / Ano-gap) is necessary as it has been shown that only after this calculation parametrical testing (e.g. by t-tests) is permissible (Schilling et al., 2017). A significant decrease (p<0.05; Bonferroni corrected) of the GPIAS-induced change in the ASR amplitude (i.e. PPI) after the trauma (PPIpost) relative to conditions before the trauma (PPIpre) in that test was rated as an indication of a tinnitus percept at that specific frequency. The effect size of the log-normalized PPI change (log(PPIpost) - log(PPIpre)) can be interpreted as a correlate of the subjective tinnitus loudness. With this behavioral approach, we were able to group animals independent of any other measurement into those with behavioural signs of a tinnitus percept and those without (Tziridis et al., 2021). The resulting distributions were tested by a chi²-test for multiple groups.

The threshold determination was performed by an automated objective approach using the root mean square (RMS) values of the ABR amplitudes fitted with a hard-sigmoid function using the background activity as offset (Schilling et al., 2019). The mean threshold was set at the level of slope change of this hard-sigmoid fit independently for each frequency and for each time point. Hearing loss (HL) was calculated by subtracting the post-trauma from the pre-trauma thresholds and tested by parametric tests. Positive HL values indicate worse hearing after the trauma; negative values indicate better hearing. The HL was calculated separately for each ear, i.e., control and trauma ear.

#### Immunohistology

Whole-mount immunohistology was performed on both cochleae of the animals (Tziridis et al., 2021). After obtaining all behavioural and electrophysiological data, the animals were sacrificed, and the cochleae of the traumatized and non-traumatized (control) ears were extracted and fixed in 4% formaldehyde for 1 hour. After de-calcification in 0.1 M EDTA for 1-2 days, the cochlear turns were usually cut into 3 pieces and immunostained for synaptic ribbon protein carboxy-terminal binding protein 2 (CTBP2). Cochlear whole-mounts were immunostained overnight at 4°C with primary antibody against CTBP2 (mouse anti-CTBP2 at 1:200; BD Transduction Labs, Heidelberg, Germany). Secondary antibody (donkey anti-mouse conjugated with Cy3, 1:400, Dianova, Hamburg, Germany) was applied for 1h at room temperature. The whole-mounts were then prepared for microscopy and the ribbon-synapses of the inner hair cells (IHC) were counted over the whole organ of corti with IHC-batches of 15 cells for averaging.

The number of synapses per IHC was counted from the microscopic images by one investigator blinded to the status of the cochlea (traumatized / control ear) or any other attribute of the animal. The continuous frequency map of the cochlea from approximately 100 Hz to roughly 40 kHz was created with the software Keyence BZ-II-Analyzer by measuring the relative distances beginning from the apex of the cochlea along the spiral, according to the published cochlear frequency map for the Mongolian gerbil (Müller, 1996). Full-focus microscopic images in the range of the synapses of the IHCs from the subjacent inner spiral bundle to the nerve terminal in the supra-nuclear region including all visible synaptic ribbons were obtained using a fluorescent microscope BZ9000 (Keyence, Neu-Isenburg, Germany) with a 40x objective (0.6 numerical aperture) and visualized using an ImageJ plugin. Labelled synapses of the IHCs were counted for groups of 15 IHC, averaged and mapped to the matching frequency. Along the frequency map, approximate octave steps were used to further group the batches of averaged synapse values for further analysis. The loss of ribbon synapses at IHC over the complete cochlear frequency map could be investigated only in a subset of mitragynine-treated animals (acute: 11 of 20; early chronic: 15 of 23) due to technical problems during histology (e.g., only one cochlea could be analysed).

### Statistical analyses

#### AUD

All quantitative data were expressed as mean ± SEM. Data were analysed using one-or two-way ANOVAs for repeated measures where appropriate, and pre-planned comparisons using Bonferroni-corrected Fisher’s LSD test or two-tailed t-test for single group comparisons.

#### Tinnitus

All statistical analyses were performed with Statistica 14 (TIBCO Software Inc., Palo Alto, USA). GPIAS effect was assessed by Bonferroni-corrected t-tests; significant negative values were seen as indicators for a tinnitus percept (T animals), all others were seen as non-tinnitus (NT) animals. Treatment animal groups were compared with the corresponding vehicle animals independently for both treatment conditions by Bonferroni-corrected chi²-tests. ABR threshold changes (control ear, trauma ear) were assessed by mixed design repeated measurement ANOVAs (repetition factor ear and factors stimulation frequency and substance application) independently for both treatment conditions. The obtained numbers of synapses were processed by calculating the difference between trauma and control ear for each animal for all corresponding cochlear frequency locations. The resulting ribbon synapse loss was evaluated for vehicle and treatment groups by two-way ANOVAs (factors cochlear frequency and behaviour at treatment start). A significance level of p<0.05 was used.

## Supplementary material

The supplementary material for this article is available.

## Data availability

The data that support the findings of this study are available from the corresponding author upon request.

## Acknowledgements

This project was in partial fulfilment of the requirements for obtaining the degree of “Dr. med.” for A.G., F.G., B. H., G.M.K., S.K., L.P., L.S., M.S., N.F., M.K., and L.W. The research was funded by the Deutsche Forschungsgemeinschaft (DFG) grant MU 2789/18-1 (C.P.M), and TRR 265 (C.P.M) and the Georg Forster award by the Alexander von Humboldt society (Z.H.).

## Author Contributions

Liubov S. Kalinichenko, Konstantin Tziridis, Zurina Hassan, Holger Schulze, and Christian P. Müller conceptualized the study and designed the analytic approach. Laura Emrich, Anna Glotz, Bernadette Hoermann, Gian-Marco Kersten, Salvador Kirmsse, Lisa Perschler, Lydia Scheidig, Martin Strotmann, Lissy Windhösel, Marie-Lou Boyer, and Liubov S. Kalinichenko performed the behavioural experiments in mice; Konstantin Tziridis, Natasha Foerst, and Melissa Kienberger performed the experiments in gerbils; Fabian Graßl and Markus R. Heinrich developed and produced fluorescence labelled mitragynine and Nassri Abdoush, Renato Frischknecht, and Anna Fejtova performed the labelling experiment. Christian P. Müller and Konstantin Tziridis drafted the manuscript; Kasia Radwanska, Markus R. Heinrich, Johannes Kornhuber, Darshan Singh, Nelson Chear Jeng Yeou, Zurina Hassan, and Holger Schulze revised the first draft. All authors critically revised the manuscript. Christian P. Müller and Liubov S. Kalinichenko had final responsibility for the decision to submit for publication.

