## Supplementary material for "Mitragynine reduces symptoms of alcohol use disorder (and tinnitus) in a self-medication of rodents"

### I. Mitragynine

Mitragynine was isolated from the leaves of *Mitragyna speciosa* (500 g), following a previously reported procedure with minor modifications (Chear et al., 2021). Briefly, the methanolic extract was subjected to acid–base extraction to obtain an alkaloid-enriched fraction (7 g). This fraction was subsequently purified by silica gel column chromatography using a stepwise gradient of hexane and ethyl acetate (EtOAc) with increasing solvent polarity. The fraction eluted with hexane/EtOAc (4:1) was collected isocratically, and yield 2.4 g of mitragynine. The purified mitragynine base was converted to its hydrochloride salt for spectroscopic characterization, which confirmed a purity of > 99% (Fig. 1).

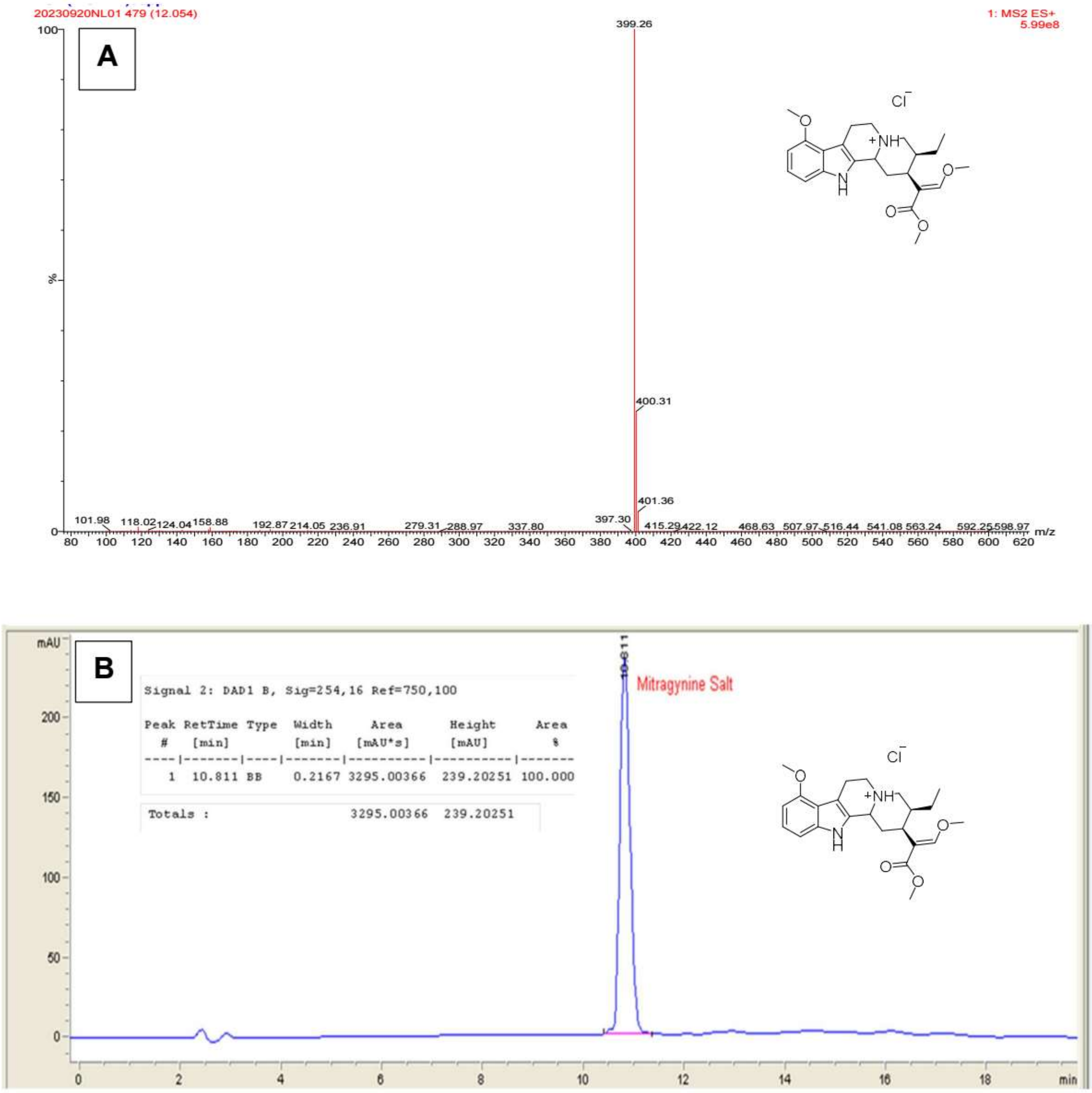


Figure 1: Purity analysis of mitragynine salt (hydrochloride): (a) Parent Ion scan using liquid-chromatography – mass spectrometer (LC-MS) ≥ 99% purity; (b) HPLC-PDA chromatogram of mitragynine salt at a wavelength of 254 nm showing ≥ 99% purity.

### II. Mitragynine-Cy5 synthesis and validation

Solvents and reagents are obtained from commercial sources and used as received. $^{1}$H-NMR and $^{13}$C-NMR spectra are recorded on the Bruker Avance 400 ($^{1}$H: 400 MHz; $^{13}$C: 101 MHz) spectrometer. Chemical shifts are referenced to residual solvent peaks in parts per million (δ): 1H δ = 7.26 ppm, 13C δ = 77.16 ppm for chloroform-*d* and 1H δ = 3.31 ppm, 13C δ = 49.00 ppm for methanol-$d_4$. Coupling constants are in Hertz (Hz). The following abbreviations are used for the description of signals: s (singlet), d (doublet), t (triplet), q (quartet), m (multiplet), bs (broad singlet) hept (heptet). Mass spectra are recorded using electron spray ionization (ESI) on a Waters Alliance HPLC system fitted with an AQUITY QDa mass detector (LRMS) or on a Bruker timsTOF pro mass spectrometer (HRMS). Silica gel (Kieselgel 60, 40-63mm, Merck) is used for flash column chromatography.

Cy5-alkine,[1] 1-azido-6-bromohexane[2] and 9-O-desmethyl mitragynine[3] were synthesized according to literature protocols.

##### 9-O-Azidohexyl mitragynine

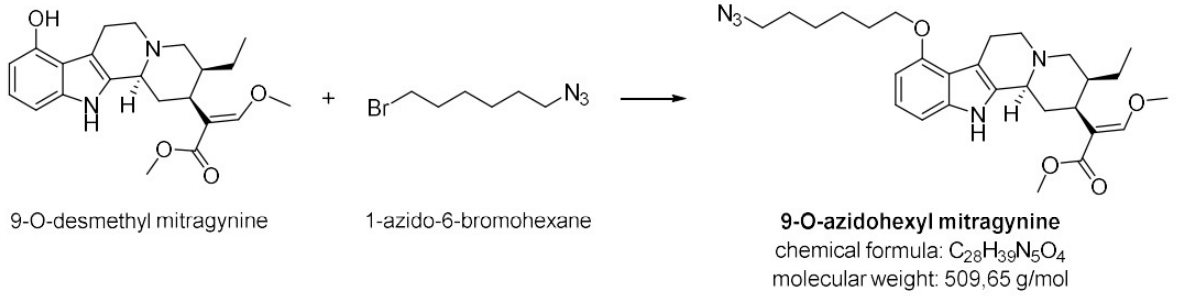


9-O-Desmethyl mitragynine (0.26 mmol, 100 mg) and potassium carbonate (1.56 mmol, 216 mg) were combined in DMF (1 mL) and 1-azido-6-bromohexane (0.34 mmol, 70 mg) dissolved in DMF (0.5 mL) was added dropwise. The mixture was stirred for 72 h at 60°C under nitrogen atmosphere. The reaction mixture was cooled to room temperature and water (20 mL) was added. The crude product was extracted with DCM (3 × 20 mL), the combined organic phases were dried over $Na_2SO_4$ and solvent was removed under reduced pressure. The crude product was purified by silica gel column chromatography (hexane:ethyl acetate 8:2 → 0:1) to obtain 9-O-azidohexyl mitragynine (57 mg, 0.11 mmol, 43 %).

$^{1}$H NMR (400 MHz, chloroform-*d*) δ 7.87 (s, 1H), 7.43 (s, 1H), 6.96 (t, J = 7.9 Hz, 1H), 6.88 (dd, J = 8.1, 0.7 Hz, 1H), 6.41 (dd, J = 7.7, 0.8 Hz, 1H), 4.09 – 3.94 (m, 2H), 3.71 (s, 3H), 3.70 (s, 3H), 3.28 (t, J = 7.0 Hz, 2H), 3.20 – 3.08 (m, 2H), 3.07 – 2.88 (m, 4H), 2.58 – 2.40 (m, 3H), 1.89 – 1.33 (m, 11H), 1.20 (t, J = 7.0 Hz, 1H), 0.89 – 0.84 (m, 3H).

$^{13}$C NMR (101 MHz, chloroform-*d*) δ 169.3, 160.7, 153.9, 137.4, 133.8, 121.9, 117.8, 111.5, 107.8, 104.1, 100.3, 67.3, 61.7, 61.4, 57.8, 53.9, 51.5, 51.5, 40.7, 40.0, 30.0, 29.4, 28.9, 26.6, 25.9, 24.1, 19.2, 12.9.

LRMS (ESI) calcd. for $C_{28}H_{40}N_5O_4$ $[M+H]^+$, 510.66, found: 510,17.

#### Mitragynine-Cy5 conjugate

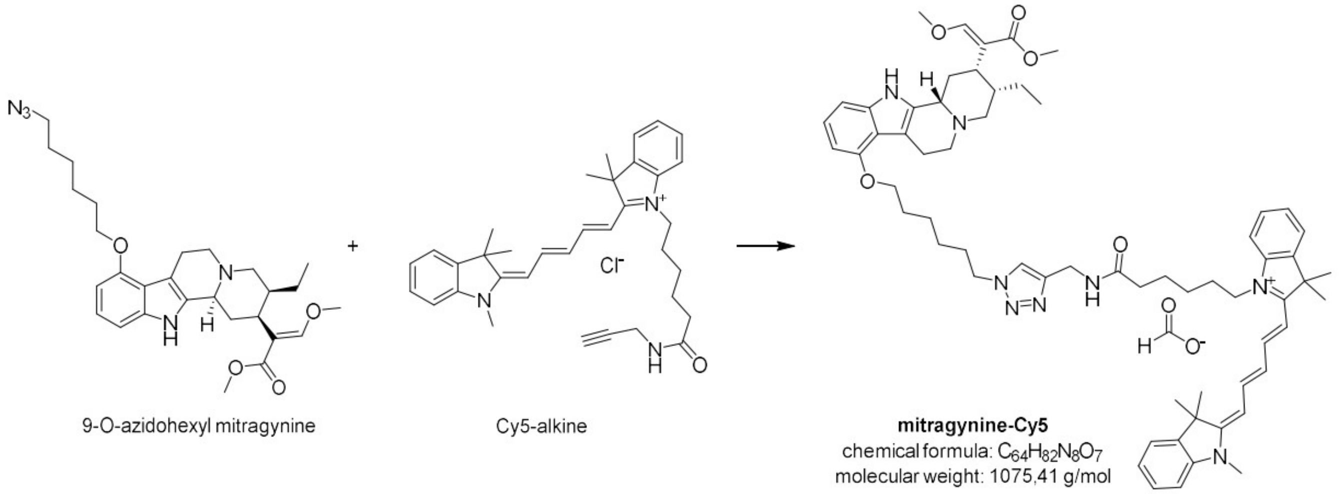

9-O-azidohexyl mitragynine; Cy5-alkine; **mitragynine-Cy5**, chemical formula: $C_{64}H_{82}N_8O_7$, molecular weight: 1075,41 g/mol

9-O-Azidohexyl mitragynine (39 µmol, 20 mg), Cy5-alkine (78 µmol, 44 mg) and copper sulfate pentahydrate (4 µmol, 1 mg) were dissolved in $H_2O$/acetonitrile (v/v = 2:1, 2 mL). Hydrazine hydrate (39 µmol, 2 µL) was added and the reaction was stirred at room temperature for 16 hours in the dark. $H_2O$ (20 mL) was added to the reaction mixture and product was extracted with DCM (3 × 20 mL). The combined organic phases were dried over $Na_2SO_4$ and the solvent was removed under reduced pressure. The crude product was purified by preparative HPLC (80:20 → 0:100 acetonitrile/water + 0.1% formic acid over 15 min, flow rate: 30 mL/min, ZORBAX Eclipse XDB-C8). Mitragynine-Cy5 was obtained as deep blue solid (13.4 mg, 32%).

$^1$H NMR (400 MHz, methanol-$d_4$) δ 8.25 (t, J = 12.8 Hz, 2H), 7.87 (s, 1H), 7.63 (s, 1H), 7.52 – 7.46 (m, 2H), 7.45 – 7.36 (m, 2H), 7.32 – 7.22 (m, 4H), 7.00 (t, J = 7.9 Hz, 1H), 6.96 – 6.89 (m, 1H), 6.64 (t, J = 12.4 Hz, 1H), 6.44 (d, J = 7.6 Hz, 1H), 6.28 (dd, J = 13.7, 4.5 Hz, 2H), 4.55 (d, J = 12.3 Hz, 1H), 4.46 – 4.30 (m, 4H), 4.12 – 3.94 (m, 4H), 3.85 (s, 3H), 3.83 – 3.78 (m, 1H), 3.76 – 3.68 (m, 4H), 3.62 (s, 3H), 3.53 – 3.34 (m, 4H), 3.29 – 3.21 (m, 1H), 2.80 – 2.70 (m, 1H), 2.54 – 2.45 (m, 1H), 2.21 (t, J = 7.5 Hz, 2H), 2.09 – 2.00 (m, 1H), 1.99 – 1.90 (m, 2H), 1.84 – 1.32 (m, 26H), 0.96 (t, J = 7.3 Hz, 3H).

$^{13}$C NMR (101 MHz, methanol-$d_4$) δ 175.7, 175.4, 174.5, 169.8, 163.3, 155.5, 155.4, 155.1, 146.3, 144.2, 143.5, 142.6, 142.5, 139.9, 129.7, 129.7, 127.5, 126.6, 126.3, 126.2, 124.5, 124.2, 123.4, 123.3, 117.7, 112.0, 111.8, 110.0, 106.8, 105.7, 104.4, 104.2, 101.4, 68.4, 64.4, 62.6, 56.8, 55.0, 52.0, 51.3, 50.5, 50.5, 44.7, 40.4, 40.4, 38.7, 36.5, 35.5, 31.5, 31.2, 30.2, 28.6, 28.1, 27.9 (2 × C), 27.8 (2 × C), 27.3, 27.1, 26.9, 26.4, 22.3, 19.4, 12.7.

HRMS (ESI) calcd. for $C_{63}H_{82}N_8O_5$ $[M+H]^{2+}$, 515.3199, found: 515.3203.

**NMR spectra**

$^{1}H$ NMR spectrum of mitragynine-Cy5 in methanol-$d_4$

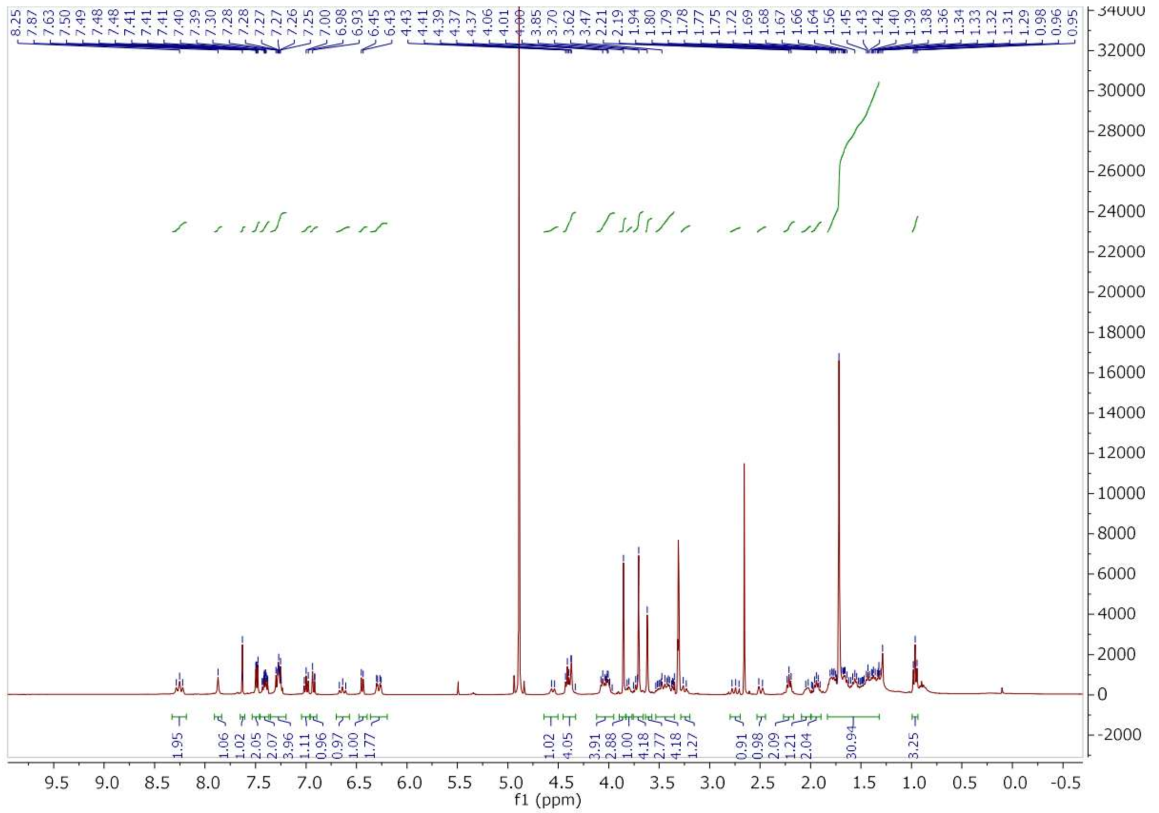

$^{13}C$ NMR spectrum of mitragynine-Cy5 in methanol-$d_4$

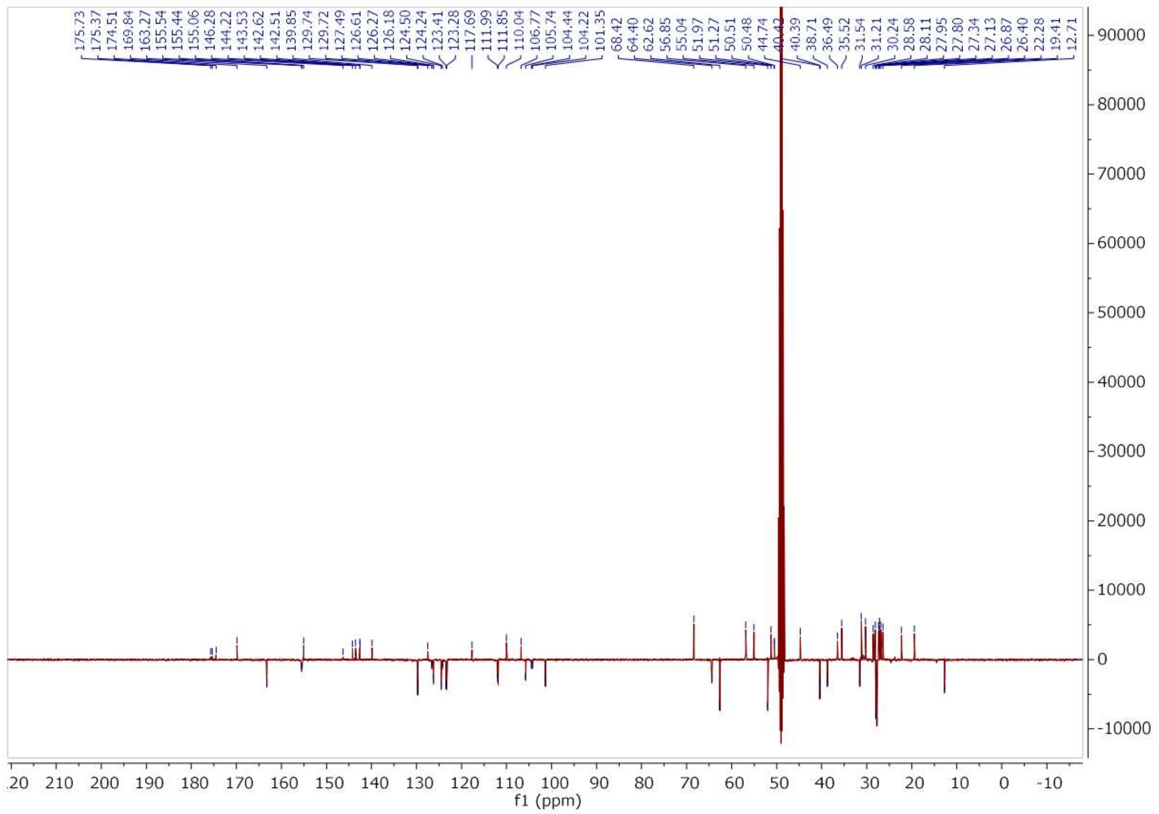

[1] O. Kaczmarek, H. A. Scheidt, A. Bunge, et al., "2′-Linking of Lipids and Other Functions to Uridine through 1,2,3-Triazoles and Membrane Anchoring of the Amphiphilic Products," *European Journal of Organic Chemistry* 2010, no. 8 (2010), https://doi.org/10.1002/ejoc.200901073.
[2] X. Cheng, G. Zhang, R. Seupel, et al., "Epoxides related to dioncoquinone B: Synthesis, activity against multiple myeloma cells, and search for the target protein," *Tetrahedron* 74, no. 38 (2018), https://doi.org/10.1016/j.tet.2018.04.056.
[3] L. C. Smith, L. Lin, C. S. Hwang, et al., "Lateral Flow Assessment and Unanticipated Toxicity of Kratom," *Chemical Research in Toxicology* 32, no. 1 (2019), https://doi.org/10.1021/acs.chemrestox.8b00218.

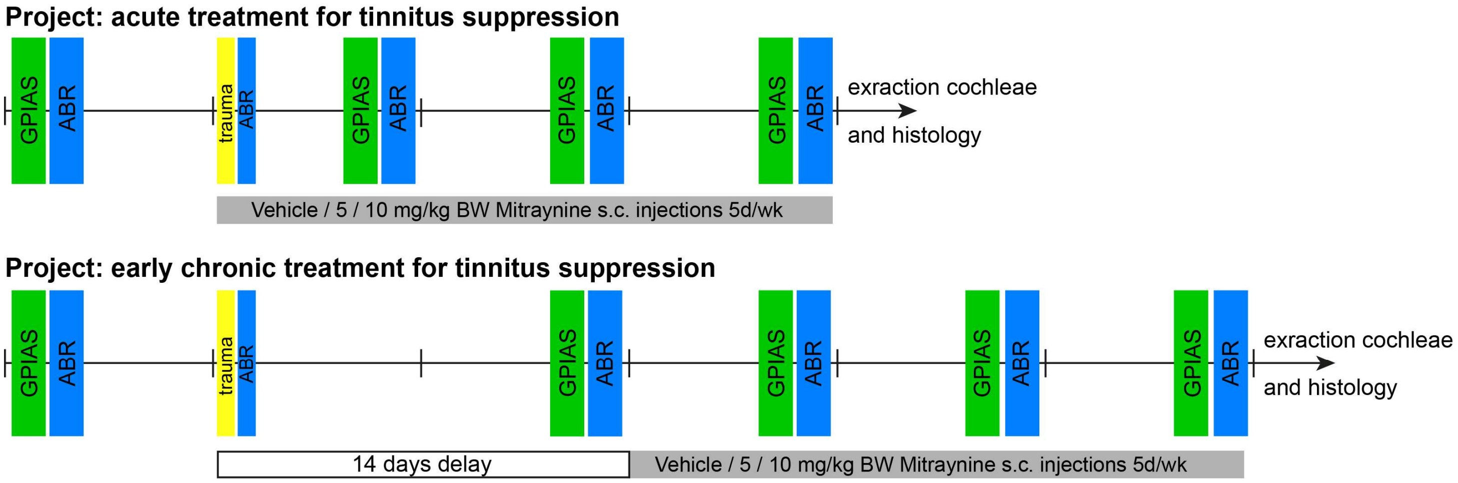


**Suppl. Figure 1** Design of tinnitus model in Mongolian gerbils.

**Suppl. Table 1:** Results of mixed repeated-measurement ANOVAS of ABR threshold shifts (i.e. hearing loss) for trauma and control ears in acutely treated animals

| ear | tested factors | F statistics | p value |
|---|---|---|---|
| trauma | group | F(2, 114)=0.42 | 0.66 |
| | frequency | F(4, 114)=9.87 | <0.001 |
| | treatment week | F(2, 228)=15.78 | <0.001 |
| | group X frequency | F(8, 114)=0.80 | 0.60 |
| | group X treatment week | F(4, 228)=7.10 | <0.001 |
| | frequency X treatment week | F(8, 228)=1.36 | 0.22 |
| | group X frequency X treatment week | F(16, 228)=1.23 | 0.24 |
| control | group | F(2, 114)=0.82 | 0.44 |
| | frequency | F(4, 114)=6.54 | <0.001 |
| | treatment week | F(2, 228)=3,71 | 0.03 |
| | group X frequency | F(8, 114)=1,14 | 0.34 |
| | group X treatment week | F(4, 228)=2,13 | 0.08 |
| | frequency X treatment week | F(8, 228)=0.79 | 0.61 |
| | group X frequency X treatment week | F(16, 228)=0.93 | 0.54 |

**Suppl. Table2:** Results of mixed repeated-measurement ANOVAs of ABR threshold shifts (i.e. hearing loss) for trauma and control ears in early chronically treated animals.

| ear | tested factors | F statistics | p value |
|---|---|---|---|
| trauma | group | F(2, 149)=0.06 | 0.94 |
| | frequency | F(4, 149)=0.63 | 0.64 |
| | treatment week | F(3, 447)=3.96 | 0.008 |
| | group X frequency | F(8, 149)=0.19 | 0.99 |
| | group X treatment week | F(6, 447)=3.77 | 0.001 |
| | frequency X treatment week | F(12, 447)=1.40 | 0.16 |
| | group X frequency X treatment week | F(24, 447)=1.00 | 0.46 |
| control | group | F(2, 143)=5.76 | 0.004 |
| | frequency | F(4, 143)=3.72 | 0.006 |
| | treatment week | F(3, 429)=2.56 | 0.054 |
| | group X frequency | F(8, 143)=0.83 | 0.58 |
| | group X treatment week | F(6, 429)=0.85 | 0.53 |
| | frequency X treatment week | F(12, 429)=0.65 | 0.80 |
| | group X frequency X treatment week | F(24, 429)=0.68 | 0.87 |
